# Characterising sensorimotor adaptation in Complex Regional Pain Syndrome

**DOI:** 10.1101/841205

**Authors:** Axel D. Vittersø, Gavin Buckingham, Antonia F. Ten Brink, Monika Halicka, Michael J. Proulx, Janet H. Bultitude

## Abstract

It has been suggested that sensorimotor conflict contributes to the maintenance of some pathological pain conditions, implying that there are problems with the adaptation processes that normally resolve such conflict. We tested whether sensorimotor adaptation is impaired in people with Complex Regional Pain Syndrome (CRPS) by characterising their adaption to lateral prismatic shifts in vision. People with unilateral upper-limb CRPS Type I (n = 17), and pain-free individuals (n = 18; matched for age, sex, and handedness) completed prism adaptation with their affected/non-dominant and non-affected/dominant arms. We examined 1) the rate at which participants compensated for the optical shift during prism exposure (i.e. strategic recalibration), 2) endpoint errors made directly after prism adaptation (sensorimotor realignment) and the retention of these errors, and 3) kinematic markers associated with strategic control. Direct comparisons between people with CRPS and controls revealed no evidence of any differences in strategic recalibration, including no evidence for differences in a kinematic marker associated with trial-by-trial changes in movement plans during prism exposure. All participants made significant endpoint errors after prism adaptation exposure, indicative of sensorimotor realignment. Overall, the magnitude of this realignment did not differ between people with CRPS and pain-free controls. However, when endpoint errors were considered separately for each hand, people with CRPS made *greater* errors (indicating more rather than less realignment) when using their affected hand than their non-affected hand. No such difference was seen in controls. Taken together, these findings provide no evidence of impaired strategic control or sensorimotor realignment in people with CRPS. In contrast, they provide some indication that there could be a greater propensity for sensorimotor realignment in the CRPS-affected arm, consistent with more flexible representations of the body and peripersonal space. Our study challenges an implicit assumption of the theory that sensorimotor conflict might underlie some pathological pain conditions.

## 1. Introduction^1^

Complex Regional Pain Syndrome (CRPS) is characterised by pain, motor deficits, and autonomic symptoms (Harden et al., 2010; Harden, Bruehl, Stanton-Hicks, & Wilson, 2007). In addition, the condition can be accompanied by neuropsychological changes (for reviews, see Halicka, Vittersø, Proulx, & Bultitude, 2020a; Kuttikat et al., 2016), including distorted representations of the body (e.g. Lewis, Kersten, McCabe, McPherson, & Blake, 2007; Moseley, 2005; Peltz, Seifert, Lanz, Müller, & Maihöfner, 2011), and altered updating of such representations (Vittersø, Buckingham, Halicka, Proulx, & Bultitude, 2020). Sensorimotor processing might also be affected by altered sensory experiences and motor deficits (Harden et al., 2010; Harden et al., 2007).

Harris (1999) proposed that incongruence between motor predictions and sensory outcomes, such as might arise in CRPS, could underlie pathological pain conditions for which there is no clear tissue damage (see also McCabe & Blake, 2007; McCabe, Blake, & Skevington, 2000). In support of this proposal, experimental manipulations that induce sensorimotor conflict have been found to increase pain and anomalous sensations in people with CRPS (Clémentine Brun, Mercier, et al., 2019) and fibromyalgia (Clémentine Brun, Mercier, et al., 2019; Martínez, Guillen, Buesa, & Azkue, 2019; McCabe, Cohen, & Blake, 2007), whereas the evidence is mixed for whiplash associated disorders (Daenen et al., 2012; Don, De Kooning, et al., 2017). Such manipulations can also increase sensory anomalies without altering pain levels in people with arthritis (Clémentine Brun, Mercier, et al., 2019), low back pain (Don et al., 2019), dancers with musculoskeletal pain (Roussel et al., 2015), and for violinists with pain (Daenen, Roussel, Cras, & Nijs, 2010; for review, see Don, Voogt, Meeus, De Kooning, & Nijs, 2017). For people with such conditions, problematic levels of sensorimotor incongruence could arise in daily life from compromised motor predictions and altered sensory feedback. Under normal circumstances, however, adaptation occurs when we are faced with conflicting information that appears to originate from the same source (e.g. in time and space; Wei & Kording, 2009). That is, the sensorimotor system can detect the discrepancy between actual and intended outcome of a movement, and adjust the spatial mappings of sensory inputs and/or the motor command for future movements to reduce the conflict (Wolpert, Diedrichsen, & Flanagan, 2011). If sensorimotor conflict underlies CRPS and related conditions, then it follows that this adaptation process could be disrupted such that the sensorimotor system is unable to compensate for incongruent information. However, the paradigms that have been used to study sensorimotor conflict and pain do not allow us to make inferences about sensorimotor adaptation.

Prism adaptation is a useful paradigm for investigating sensorimotor adaptation because it enables scrutiny of several distinct sensorimotor processes (e.g. strategic recalibration, sensorimotor realignment, and retention; Fig. 1). These processes and their cortical mechanisms have been studied in great detail (for reviews see Jacquin-Courtois et al., 2013; Panico, Rossetti, & Trojano, 2020). A typical prism adaptation procedure involves performing ballistic pointing movements while wearing goggles fitted with prismatic lenses that create a lateral optical shift (Held & Freedman, 1963; Redding, Rossetti, & Wallace, 2005; Von Helmholtz, 1924). During prism exposure, participants initially make pointing errors in the direction of the prismatic shift. These pointing errors will quickly reduce as movements are repeated, such that pointing is once again accurate within about a dozen trials. At first, the pointing errors are reduced mainly through “strategic recalibration”, a somewhat conscious process in which participants deliberately adjust their aim and/or mentally rotate the target location to correct for the visual shift (Rossetti, Koga, & Mano, 1993). In the longer term (e.g. over 50-100 movements or more) the spatial reference frames that coordinate visual, motor, and proprioceptive processing gradually realign to compensate for the optical distortion introduced by the prisms (“sensorimotor realignment”; Jeannerod & Rossetti, 1993; Redding et al., 2005). Once the prism goggles are removed, people typically make pointing errors in the direction opposite to the optical displacement (the prism adaptation “after-effect”). With further repeated pointing, these errors will quickly reduce if they are visible to the participants (i.e. the hand is not occluded). Even after an extended washout period, a small degree of the prism adaptation after-effect can still be observed when pointing movements are performed without visual feedback (retention). The retention of prism adaptation after-effects reflects the degree to which the realignment of visual and proprioceptive reference frames is maintained (Prablanc et al., 2019).

**Figure 1.**
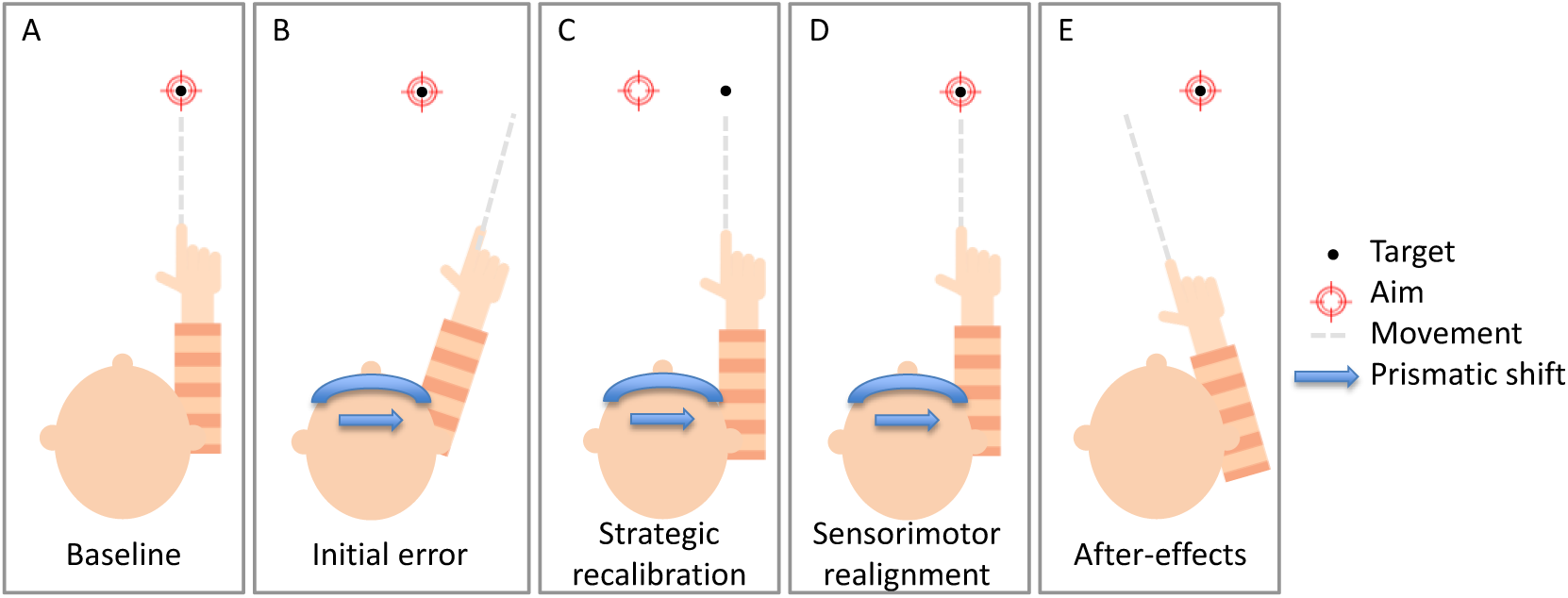
The processes involved in prism adaptation are depicted. Baseline pointing errors (A) and prism after-effects (E) are measured without wearing goggles. During early prism exposure trials (B) participants make initial errors in the direction of the prismatic shift (to the right in this example). In the first few trials of prism exposure participants correct their pointing mainly through strategic recalibration (e.g. deliberately aiming to the left of target; C). Over the longer term, sensorimotor realignment occurs (D) such that the represented visual location of the target is brought into alignment with the felt location of the arm, and participants no longer need to deliberately mis-aim to reach the target. Once the goggles are removed participants make errors in the direction opposite to the prismatic shift (i.e. after-effects; E), which are leftward in this example.

The distinct sensorimotor processes involved in prism adaptation can be quantified by measuring endpoint errors and kinematic markers, and can be expressed with exponential decay functions. Strategic recalibration (Fig. 1C) can be measured by the reduction in endpoint errors during early prism exposure. Sensorimotor realignment (Fig. 1D) and its retention are typically indexed by the magnitude of the endpoint errors made once the goggles are removed (e.g. during open-loop pointing; Fig. 1E), relative to baseline (Fig. 1A; Prablanc et al., 2019). To quantify the rapid changes during early prism exposure, endpoint errors can be fitted to an exponential decay function (Facchin, Bultitude, Mornati, Peverelli, & Daini, 2018; Martin, Keating, Goodkin, Bastian, & Thach, 1996; Nemanich & Earhart, 2015; O’Shea et al., 2014). However, analysis of endpoint error alone risks overlooking some of the sensorimotor processes involved in prism adaptation. For example, some degree of online correction is common once participants become aware of their pointing error, which can mask some of the other processes involved in strategic recalibration. One way to circumvent this problem is to examine the kinematics of the pointing movement prior to visual feedback becoming available to the participant. O’Shea et al. (2014) found that participants updated their aim to compensate for the error made on a previous trial. Specifically, the direction in which participants initiated a movement (i.e. the angle of the tangential velocity vector at peak acceleration) was adjusted to compensate for the endpoint error made on the previous trial (e.g. aiming more towards the left after making a rightwards error). Therefore, kinematic recordings of arm movements during prism exposure allow for the trial-by-trial changes in movement plans to be computed independent of any online control, enabling insights into the process of strategic recalibration.

We aimed to investigate sensorimotor adaptation in pathological pain by characterising prism adaptation in people with CRPS affecting one upper-limb relative to pain-free controls. We investigated the development and extent of strategic recalibration; and the development, magnitude, and retention of sensorimotor realignment. Participants underwent prism adaptation once with each hand, which enabled us to compare outcomes between Groups (CRPS, controls), and the Hand used (affected/non-dominant, non-affected/dominant).

Table 1 summarises the hypotheses of the study and the patterns of findings that would support these hypotheses. Based on the sensorimotor theory of pain, we predicted that people with CRPS would have problems compensating for incongruent sensory and motor information, showing impaired strategic recalibration (Hypothesis 1) and sensorimotor realignment (Hypothesis 2) based on endpoint errors. We expected that compared to pain-free controls, people with CRPS would require more trials for endpoint errors to asymptote during closed-loop prism exposure trials, that they would show smaller magnitudes and less retention of after-effects during open-loop trials, and greater residual error. We also hypothesised that the sensorimotor realignment might develop and/or decline at different rates for people with CRPS compared to controls based on open-loop endpoint errors (Hypothesis 3). We hypothesized that people with CRPS would show less evidence of trial-by-trial changes in movement plans to compensate for the prismatic shift than controls, as measured from kinematic analysis during closed-loop trials (Hypothesis 4). The trial-by-trial changes allow for deliberate changes in movement plans to be measured, which are an important strategy involved in strategic calibration (O’Shea et al., 2014), whilst eliminating any contribution from online corrections. As certain deficits (e.g. proprioceptive; Bank, Peper, Marinus, Beek, & van Hilten, 2013) can be apparent in the healthy limb, although more subtle than in the CRPS-affected limb, we expected that people with CRPS could show differences to controls in both arms. However, we also considered that any deficits in strategic control, sensorimotor realignment, and/or trial-by-trial changes in movement would be either limited to, or more apparent in, the CRPS-affected arm compared to the non-affected arm (Hypothesis 5).

**Table 1.** Outline of hypotheses and the patterns of data that would support these hypotheses.

| Hypothesis | Evidenced by | Results section |
| --- | --- | --- |
| **1) People with CRPS will show impaired strategic recalibration relative to controls** |  |  |
| *Primary outcome* | The number of closed-loop trials needed to compensate for initial errors during prism exposure (i.e. $1/b$; Fig 1C) will be larger for people with CRPS than pain-free controls | 3.2.1 |
| *Secondary outcome* | The number of closed-loop trials needed to compensate for initial errors during washout trials (i.e. $1/b$) will be larger for people with CRPS than pain-free controls | 3.2.2 |
| **2) People with CRPS will show impaired sensorimotor realignment relative to controls** |  |  |
| *Primary outcome* | The magnitude of pointing errors made on open-loop trials after prism exposure and retention trials, which should be in the direction opposite to the prismatic shift (Fig. 1E), will be smaller for people with CRPS than pain-free controls | 3.3.1 |
| *Secondary outcomes* | The residual error from closed-loop pointing movements during prism exposure and retention trials (i.e. $c$) will be larger for people with CRPS than pain-free controls | 3.2.1,<br>3.2.2 |
| **3) Sensorimotor realignment might develop and/or decline at different rates for people with CRPS compared to controls** |  |  |
|  | The magnitude of pointing errors made on interim open-loop trials during prism exposure phase (development) and during washout phase (decline) will differ between people with CRPS than pain-free controls | 3.3.2,<br>3.3.3, |
| **4) People with CRPS would show less evidence of trial-by-trial changes in movement plans than controls** |  |  |
|  | Kinematic markers of trial-by-trial changes during closed-loop trials will be less evident in people with CRPS than in pain-free controls | 3.4. |
| **5) Any deficits in strategic control, sensorimotor realignment, and/or trial-by-trial changes in movement would be either limited to, or more apparent in, the CRPS-affected arm compared to the non-affected arm** |  |  |
|  | Differences found in any of the analyses addressing hypotheses 1-4 will depend on the Hand being used for people with CRPS | 3.2.-3.4. |

## 2. Materials and methods

We used a single-session mixed design in which participants with CRPS and controls each underwent prism adaptation using each arm in the same session, and we compared the performance between the two arms and between groups. We report how we determined our sample size, all data exclusions, all inclusion/exclusion criteria, whether inclusion/exclusion criteria were established prior to data analysis, all manipulations, and all measures in the study. The study was preregistered on the Open Science Framework (https://osf.io/6jpfg/).

### 2.1 Participants

Seventeen people with CRPS type 1 predominantly affecting one upper limb (*M*_age_ = 53.53 years, *SD* = 11.67; 16 female; 14 right-handed; 9 left-affected CRPS; Table 2) were recruited through the UK national CRPS registry and from our own database. The latter is an internal database of people with CRPS consenting to be contacted about research who have been referred to us from the Royal United Hospitals (Bath, UK), Oxford University Hospitals (Oxford, UK), and Royal National Orthopaedic Hospital (London, UK) NHS Foundation Trusts; or who have contacted us directly. We decided on our sample size pragmatically, based on the maximum number of people with CRPS we could feasibly recruit and test given financial and time constraints. Twelve participants met the Budapest research criteria for CRPS (Harden et al., 2010; Harden et al., 2007), three met the clinical criteria, and two met the criteria for CRPS not otherwise specified. Fourteen of the people with CRPS had previously participated in a randomized control trial of prism adaptation for pain relief in which half of the participants underwent two weeks of twice daily prism adaptation treatment using their affected hand and half performed identical movements while wearing goggles fitted with neutral lenses (Halicka, Vittersø, et al., 2020b; Halicka, Vittersø, Proulx, & Bultitude, 2020b). There was an average of 17.13 months (*SD* = 5.69, min = 7) between participants completing the two-week exposure phase of the randomized control trial and when they took part in the current study. Eight of these participants received prism adaptation treatment, and six received sham treatment. The remaining participants with CRPS had never undergone prism adaptation before. Eighteen pain-free control participants (*M*_age_ = 54.17 years, *SD* = 12.22; 17 female; 15 right-handed) who were matched to the participants with CRPS for age (±5 years), sex, and self-reported handedness were recruited from a community sample. None of the pain-free control participants had undergone prism adaptation before.

**Table 2.**
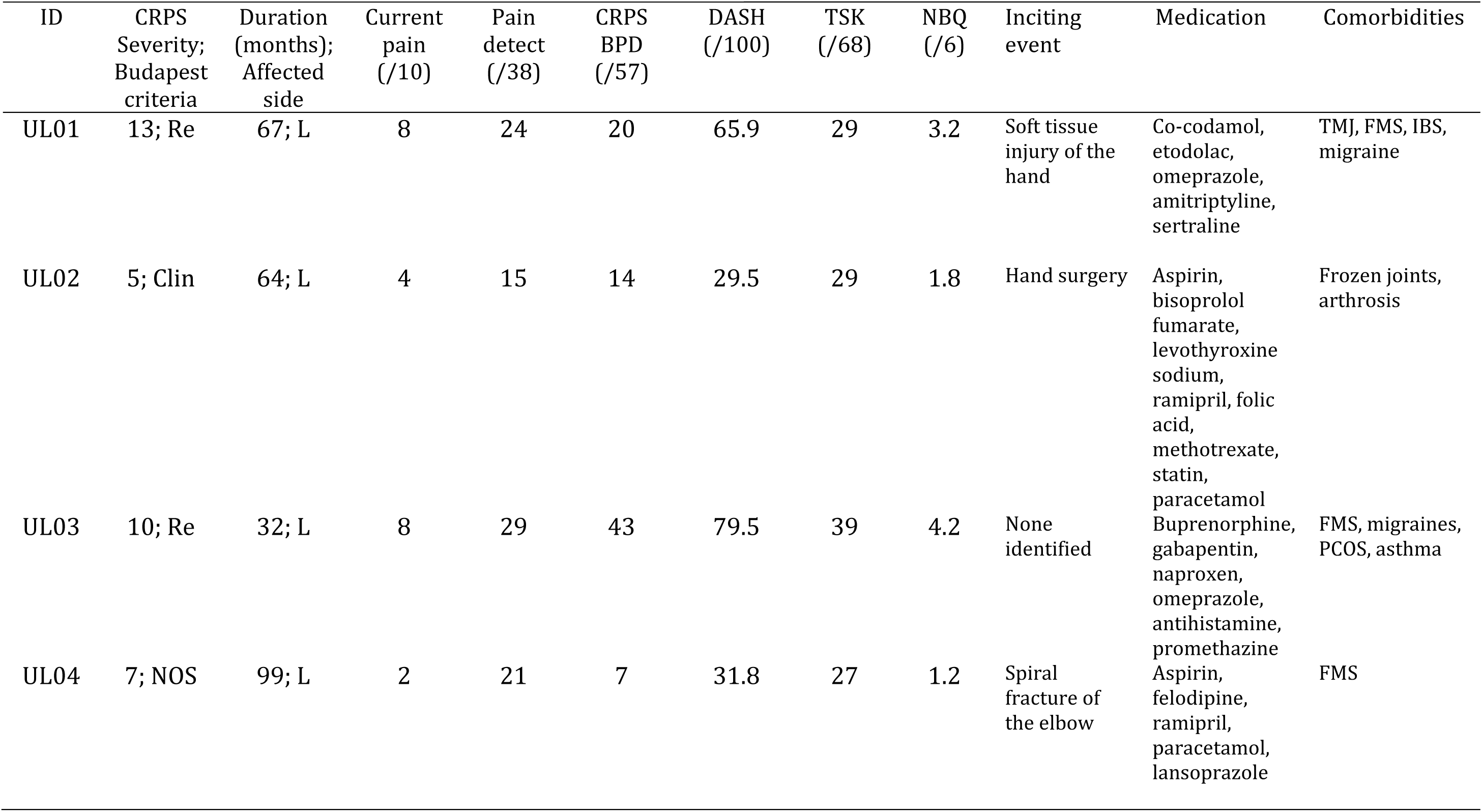
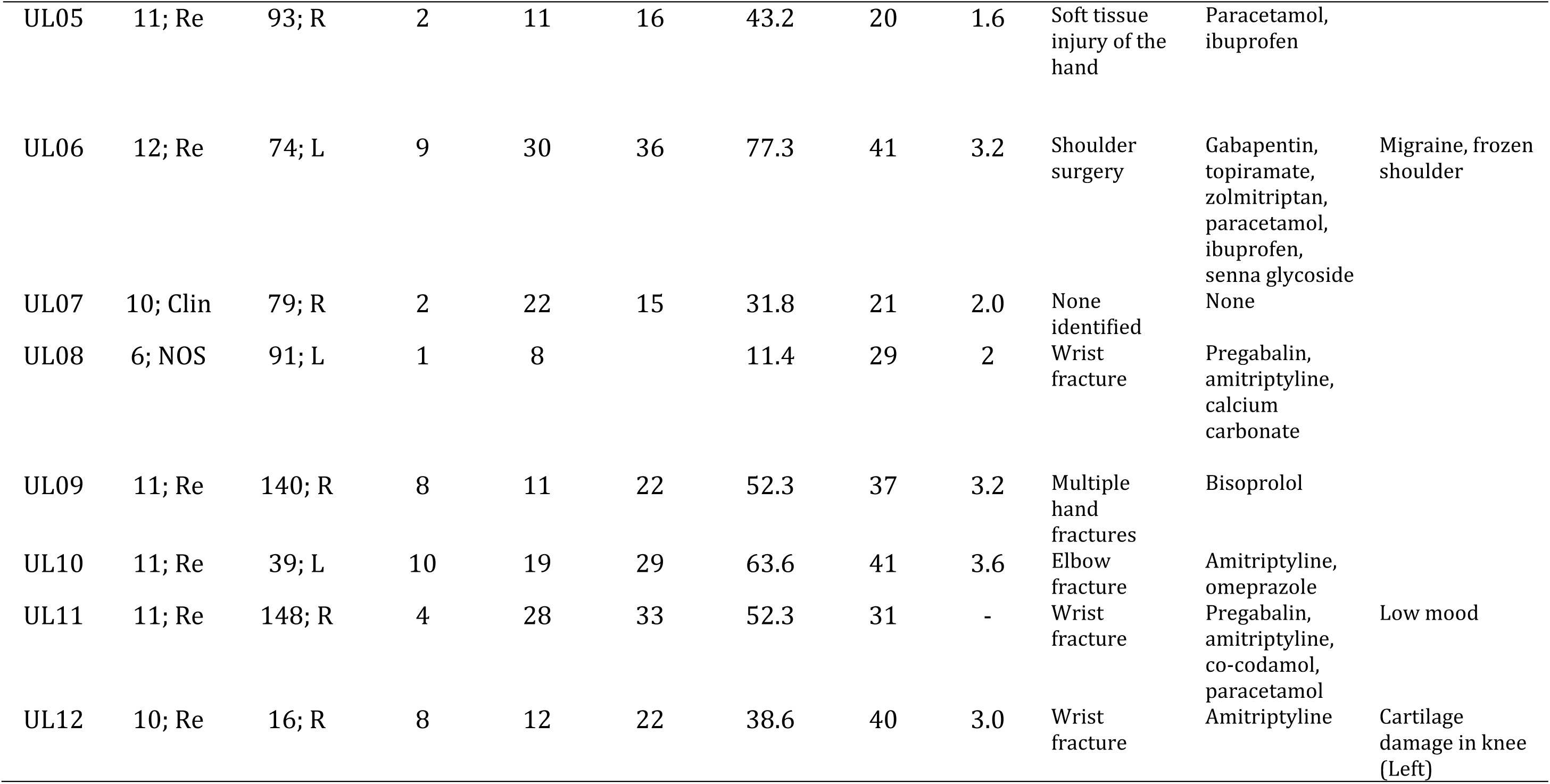
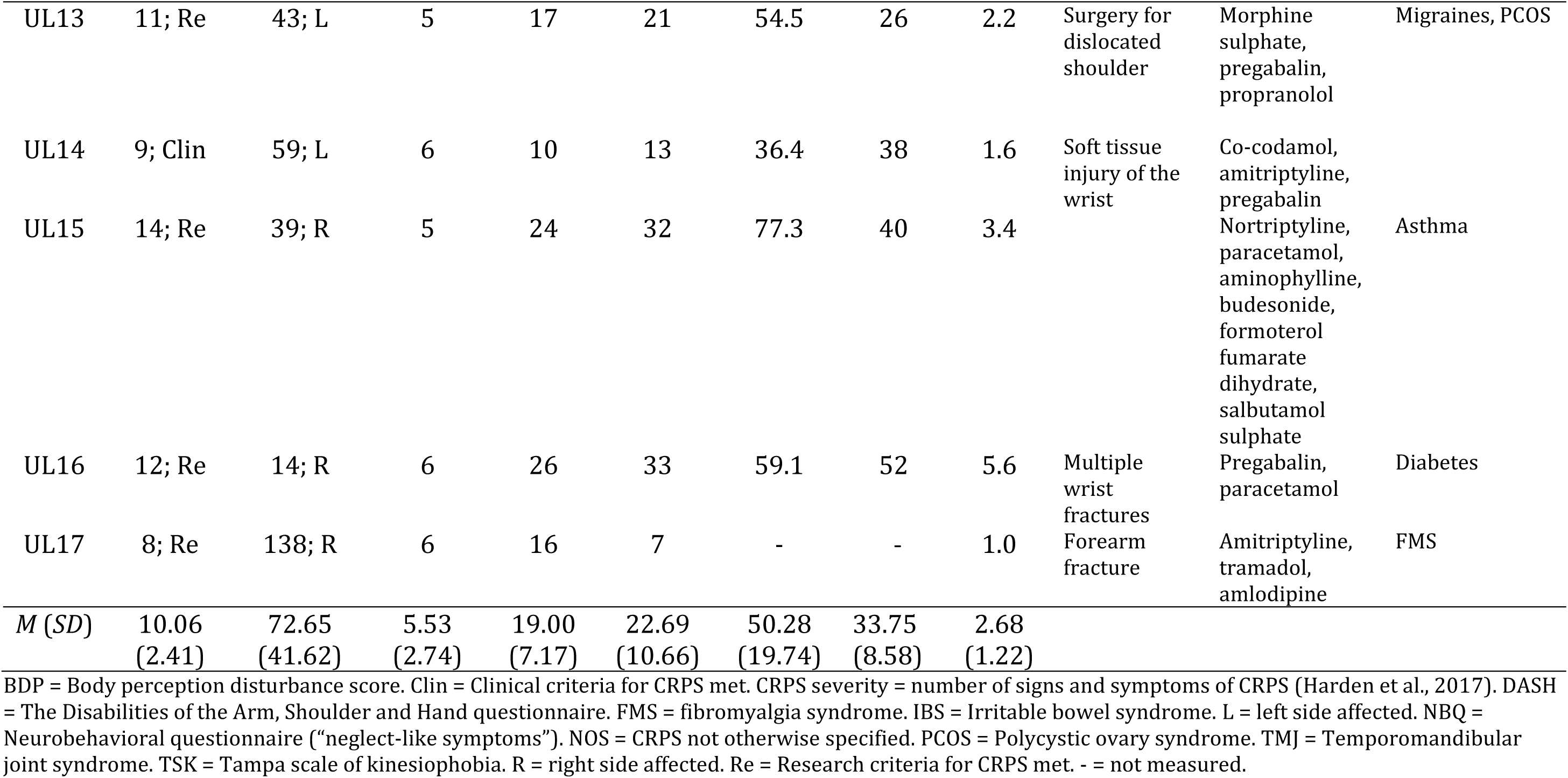
Clinical information for people with upper limb CRPS.

Participants were excluded if they reported a history of brain injury, brain disorders, or psychiatric disorders that can be associated with pronounced perceptual changes (e.g. schizophrenia; Tseng et al., 2015). Because the study involved exposure to a magnetic motion capture system, we also excluded people with a pacemaker, spinal cord stimulator or similar devices, and women who were pregnant or breastfeeding. All participants reported having normal or corrected-to-normal vision, and sufficient motor abilities to perform the movements required for the task. The study complied with the 2013 declaration of Helsinki and had ethical permission from the UK Health Research Authority (REC reference 12/SC/0557).

### 2.2 Stimuli and procedure

#### 2.2.1 Questionnaire measures

After providing informed written consent, participants completed questionnaire measures. All participants completed the Edinburgh handedness inventory (Oldfield, 1971), in which a negative score (<-40) indicates left-handedness, and a positive score (>40) indicates right-handedness. Three people with CRPS were classed as left-handed, four as ambidextrous, and eight as right-handed. Two control participants were classed as left-handed, three as ambidextrous, and 11 as right-handed.

People with CRPS completed additional questionnaires as follows (Table 2). Neuropathic components of pain were assessed by the pain DETECT questionnaire (Freynhagen, Baron, Gockel, & Tölle, 2006), where a score above 19/38 suggests that a neuropathic component is likely (>90% probability). The QuickDASH (Gummesson, Atroshi, & Ekdahl, 2003) was used to evaluate the degree of upper limb disability, where more severe disability is indicated by a higher score (/100). The Tampa Scale of Kinesiophobia (Miller, Kori, & Todd, 1991) was used to measure pain-related fear of movement and re-injury, where scores range from 17 (no kinesiophobia) to 68 (highest possible kinesiophobia). Body representation distortion was assessed by the Bath CRPS Body Perception Disturbance Scale (Lewis & McCabe, 2010), scored from zero (no body perception disturbance) to 57 (highest possible body perception disturbance).

Finally, severity of “neglect-like symptoms” was assessed by the Neurobehavioral questionnaire (Frettlöh, Hüppe, & Maier, 2006; Galer & Jensen, 1999), scored from one (no “neglect-like symptoms”) to six (highest possible severity of “neglect-like symptoms”).

#### 2.2.2 Prism adaptation

Participants performed a dynamic prism adaptation paradigm (Prablanc et al., 2019) that involved both open- and closed-loop trials (Fig. 2). Prism adaptation is appropriate for testing sensorimotor adaptation as it is thought to favour context-independent sensorimotor processes, and thus offers greater insight into the workings of the sensorimotor system than other paradigms that rely more on context-dependent processes (e.g. visuo-motor rotation; for review, see Fleury, Prablanc, & Priot, 2019). Because there has been some suggestion that adaptation to optical shifts towards the affected side might exacerbate pain for people with CRPS (Sumitani, Rossetti, et al., 2007), we used adaptation to optical shifts away from the affected/non-dominant side (leftwards for 9/17 people with CRPS; leftwards for 3/18 controls). The choice of prismatic shift was therefore consistent with previous studies suggesting a therapeutic effect of prism adaptation away from the CRPS-affected limb (Bultitude & Rafal, 2010; Christophe et al., 2016; Sumitani, Rossetti, et al., 2007). From a sensorimotor perspective, the magnitude of the sensorimotor realignment is not significantly different for leftward shifting and rightward shifting prisms (Schintu et al., 2017). As we were primarily interested in testing the processes involved in sensorimotor adaptation, rather than pain in response to conflicting sensorimotor information, we did not consider there to be any benefit to using lenses with that induced an optical shift towards the affected side. For the prism exposure trials, participants wore goggles fitted with Fresnel lenses that shifted vision laterally by 35 dioptres (∼19°). Participants completed the prism adaptation protocol with each hand in a randomised and counterbalanced order. During open-loop trials participants’ vision of their hand was occluded, and they performed pointing movements to a central visual target (i.e. 0°). For closed-loop trials, participants had vision of their hand when their arm was fully extended, whereas the arm from the wrist up was concealed (i.e. terminal exposure). Visual targets for closed-loop trials were 10° to the left or right of centre. For each hand, participants completed closed-loop and open-loop trials across five phases (see Fig. 2). First, in the baseline phase, participants completed one block of 20 closed-loop trials and one block of 15 open-loop trials with unperturbed vision. Second, in the prism exposure phase, they performed 100 closed-loop trials towards targets viewed through the prism goggles, split into six blocks (Prism exposure – closed loop 1 [P-CL1], P-CL2, P-CL3, P-CL4, P-CL5, P-CL6). The blocks of closed-loop trials were separated by blocks of two open-loop trials (Prism exposure – open loop 1 [P-OL1], P-OL2, P-OL3, P-OL4, P-OL5) towards targets view through unperturbed vision. Third, in the after-effect phase, participants performed another block of 15 open-loop trials towards targets viewed with unperturbed vision. Fourth, in the washout phase they performed 60 closed-loop trials towards targets viewed with unperturbed vision, split into six blocks (Washout – closed loop 1 [W-CL1], W-CL2, W-CL3, W-CL4, W-CL5, W-CL6). The blocks of closed-loops trials were once again separated by blocks of two open-loop trials (Washout – open loop 1 [W-OL1], W-CL2, W-OL3, W-OL4, W-OL5) towards targets view through unperturbed vision. Finally, in the retention phase participants performed one block of 15 open-loop trials with unperturbed vision. The entire procedure was completed once for each hand, in a counterbalanced order. For each set of 10 closed-loop trials, five left targets and five right targets were presented, in a randomised order.

**Figure 2.**
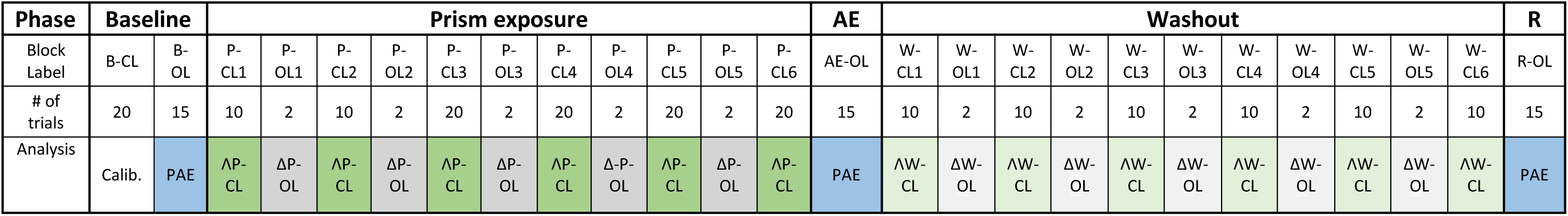
Prism adaptation protocol. Colours indicate what analysis each phase or block corresponds to. AE = after-effect; B = baseline; Calib. = calibration trials; CL = closed-loop pointing; OL = open-loop pointing; PAE = prism adaptation after-effects; P-CL = closed-loop trials during prism exposure phase; P-OL = open-loop trials during prism exposure phase; R = retention of after-effect; W-CL = closed-loop trials during washout phase; W-OL = open-loop trials during washout phase; Δ = change in endpoint errors during open-loop trials; Δ = exponential decay of endpoint errors during closed-loop trials.

The first four people with CRPS to participate performed an additional 42 trials with each hand (two open-loop trials followed by 40 closed-loop trials) at the end of the washout phase. Despite their relatively good upper limb mobility, they found it difficult to complete all 287 pointing movements with their affected hand. We therefore reduced the number of trials in the washout phase for the remaining participants to that represented in Fig. 2, and updated the preregistration accordingly (https://osf.io/6jpfg/). This resulted in a total of 245 trials per hand.

Participants were seated at a custom-built non-ferrous table (Fig. 3) with a Velcro-adjustable chin-rest affixed to the edge closest to the participant, and the magnet from the motion capture system attached to the underside of the table at the edge furthest away from the participant. A motion tracking sensor (7.9 mm × 7.9 mm × 19.8 mm) was placed on each of the participants’ index fingers with medical tape. To start a trial, participants placed their index finger on a raised tactile point (∼1 cm diameter) near to the trunk that was aligned with their body midline and the central target (the “start location”). The trial was programmed to start only if one of the sensors was first detected within ±2 cm laterally and ±3 cm distally of the start location. If a sensor was detected, a red light-emitting diode (LED) appeared in one of the target locations after a pause of 1 s. After a further 1 s, an audio cue (200 ms, 800 Hz) sounded. Participants were instructed that upon hearing the audio cue, they should point as quickly as possible with their fully extended arm and place their finger on the line leading to the visual target (Fig. 3). The visual target stayed illuminated for a further 3 s after the onset of the audio cue. Participants were instructed to bring their hand back to the start location upon the target extinguishing. The experimenter then pressed a computer key to allow the script to proceed to the next trial, which would once again start only if one of the sensors was detected within ±2 cm x ±3 cm of the start location. If interim open-loop trials (e.g. P-OL1, W-OL2) were completed incorrectly (e.g. a false start), they were repeated. No other trials were repeated.

**Figure 3.**
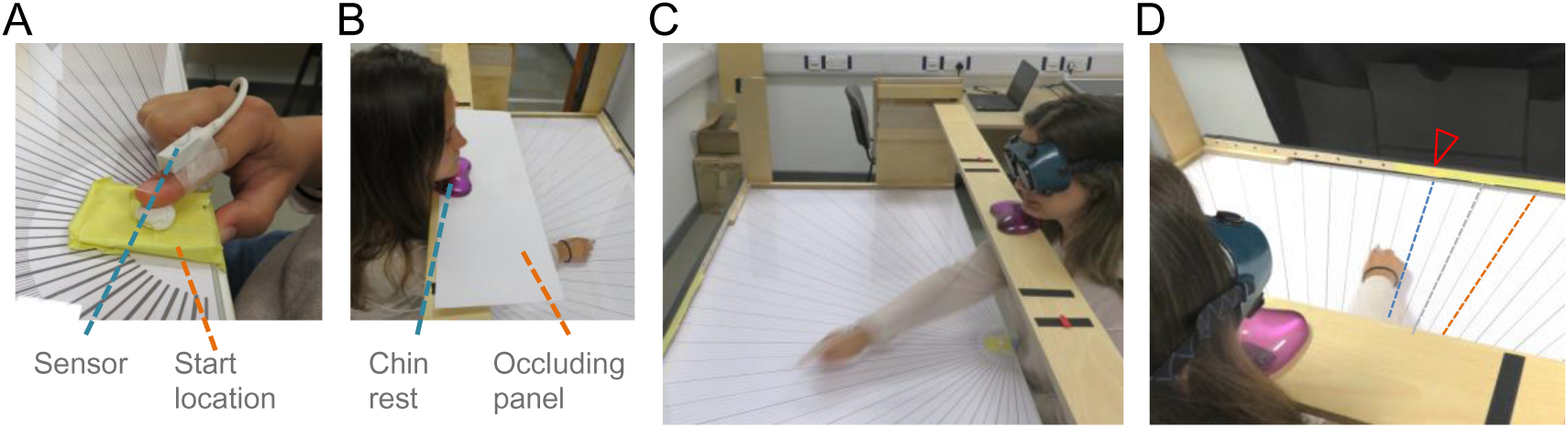
The prism adaptation table, chinrest, sensor (A), start location (A), and occluding panel (B). Panel B depicts an open-loop trial. Panels C and D depict closed-loop trials. The red arrow (D) indicates an example target location, with 10° left (blue), 10° right (orange), and central (i.e. 0°; grey) target axes superimposed.

We recorded 3D kinematic data at 240 Hz using an electromagnetic motion capture system (trakSTAR™, 3D Guidance^®^, Northern Digital Incorporated). The kinematic data was low-pass filtered using a second-order dual-pass Butterworth filter at 10 Hz. We calculated instantaneous velocities and accelerations by differentiating the data with a 5-point central finite difference algorithm, twice per axis. Velocity vectors were combined to yield resultant velocity, which was used to determine movement onset, and movement offset. The threshold for movement onset was set at 50 mm/s. Because people with CRPS can have motor impairments (e.g. spasms or arrests), we considered multiple criteria for movement offset for all participants: a threshold of 50 mm/s, and the point at which the sensor returned to the same vertical location as movement onset. The latter was taken as proxy of the hand being placed on the table. We visually inspected all trials (displacement, and resultant velocity plots), and manually adjusted movement onsets and movement offsets to satisfy the above criteria when needed. We deleted trials where a false start was detected (i.e. when movements faster than 50 mm/s were detected for the first sample; 2.52% of trials), and rotated movement trajectories to correct for a calibration error.

### 2.3 Statistical analyses and inference criteria

#### 2.3.1 Endpoint errors

Our primary analyses were of endpoint errors from closed-loop (3.2) and open-loop (3.3) trials. We calculated endpoint errors (°) as the angle between a two-dimensional straight line connecting the start location and movement offset, and a straight line from the start location to the target (i.e. the target axis). Endpoint errors made towards the affected/non-dominant side were expressed as negative values. We analysed trial level data using a linear mixed model regression, with participant ID and trial number as random effects. However, we did not include trial as a random effect for analyses where there was only one datapoint for each level of the fixed effect (e.g. 1/*b*, or *c*).

#### 2.3.2 Exponential decay

For closed-loop trials during the prism exposure phase (3.2.1; P-CL1 – P-CL6), and the washout phase (3.2.2; W-CL1 – W-CL 6) we fitted exponential decay functions (*x* = *a* × e^-*b* × *n*^ + *c*) to endpoint errors for each person and each Hand (Facchin et al., 2018; Martin et al., 1996; Nemanich & Earhart, 2015; O’Shea et al., 2014). We considered *x* as the endpoint error; *a* the initial error; *b* the decay constant; *c* the residual error; and *n* the trial number. The rate of error correction (i.e. the decay factor) was expressed as the inverse of *b* (i.e. 1/*b*). The inverse of *b* equates to the half-life of the endpoint error reduction curve, and therefore indicates half the number of trials needed for endpoint errors to reach the asymptote (i.e. the residual error *c*).

To examine whether people with CRPS would show impaired strategic control relative to controls (Hypothesis 1), we compared the decay factor (i.e. 1/*b*) between Groups and the Hand used. We analysed the residual error (i.e. *c*) to address Hypothesis 2, also comparing between Groups and the Hand used. To perform these analyses, we used linear mixed models regressions with participant ID as a random effect.

However, our primary analysis to address Hypothesis 2 (that people with CRPS would show impaired sensorimotor adaptation relative to controls) was to compare “raw” open-loop endpoint errors (2.3.1) across the different experimental phases (Fig. 2). That is, we analysed endpoint errors from open-loop trials without fitting an exponential decay function. We compared endpoint errors between Groups, Hand, and Phase (baseline, after-effect, retention). There were 15 open-loop pointing trials per phase for each hand.

To address Hypothesis 3, that sensorimotor realignment might develop and/or decline at a different rate for people with CRPS, compared to controls, we examined the development of the prism adaptation after-effects during the prism exposure phase (3.3.2), and their decline during the washout phase (3.3.3). As with the primary analysis to address Hypothesis 2, we analysed endpoint errors without fitting an exponential decay function for these analyses. We performed separate analyses for the open-loop blocks during the prism exposure and the washout phases (Fig. 2). For the prism adaptation phase, we compared endpoint errors between Groups, Hand, and PA Block (P-OL1 – P-OL5); for the washout phase, we compared endpoint errors between Groups, Hand, and Washout Block (W-OL1 – W-OL5).

#### 2.3.3 Trajectory orientations

We derived kinematic markers (3.4) that have previously been associated with strategic recalibration (3.4.1) and sensorimotor realignment (3.4.2) during prism exposure (O’Shea et al., 2014). Specifically, we computed the tangential velocity vectors for peak acceleration (initial trajectory orientation) and peak deceleration (terminal trajectory orientation) and expressed them as the angle (°) relative to the target axis.

To address Hypothesis 4, that people with CRPS would show less evidence of trial-by-trial changes in movement plans to compensate for the prismatic shift than controls, we analysed the trial-by-trial change in initial trajectory orientations during early trials (i.e. closed loop trials 1 to 10 during prism exposure; P-CL1) for each Group, and Hand. These trial-by-trial changes allow for a specific strategy involved in strategic calibration to be examined (O’Shea et al., 2014), whilst eliminating any contribution from online corrections. We correlated the magnitude of endpoint errors on a given trial (*n*) during early prism exposure with the change in initial trajectory orientation on the subsequent trial (*n*+1). We performed this correlation on detrended data for only the early prism exposure trials because error correction is typically only evident in these. To detrend the data, endpoint errors and initial trajectory orientations were fitted to the same exponential decay function as described in sections 2.3.2 (i.e. *x* = *a* × e^-*b* × *n*^ + *c*), and then the residuals were computed by subtracting the predicted values (i.e. *x*) from the observed values for each trial. The *t*-value for the correlation between these variables was calculated for each participant and each hand. If endpoint errors and initial trajectory orientations are unrelated then the *t*-values from these individual correlations should have a Gaussian distribution centred around zero. We aimed to test whether, for each Group and Hand, there was a linear relationship between endpoint errors on a given trial (*n*) and the change in movement plan on the next trial (*n*+1) by using one-sample *t*-tests to compare the individual participant *t*-values to zero. This analysis has been used previously to examine the changes in feedforward motor control during early prism exposure (O’Shea et al., 2014).

To further address Hypothesis 4, we performed unilateral Kolmogorov-Smirnov distribution tests (Vindras, Desmurget, & Baraduc, 2012) on the *p*-values obtained from individual correlations to see if they were biased towards zero. If individual *p*-values are biased towards zero it would provide further support for the presence of a linear relationship between endpoint errors on a given trial and the updated movement plan of the subsequent trial. This analysis therefore sheds light on the process of strategic error reduction during early prism exposure (O’Shea et al., 2014).

In our pre-registration we aimed to compare the terminal trajectory orientations between Groups, however this was not possible. Many participants made corrective finger movements in the later stage of a movement, once they became aware that they were about to miss the target. These late finger movements limited the information that could be derived from the point of peak deceleration (e.g. terminal trajectory orientations). When we filtered the data to remove these corrective finger movements the sample sizes were too small to make meaningful comparisons between Groups (CRPS n_affected_ = 8; CRPS n_non-affected_ =14; control n_non-dominant_ = 10; controls n_dominant_ = 15). Therefore, we do not report on the analyses of terminal trajectory orientations.

#### 2.3.4 Inference criteria

We processed and analysed the data in MATLAB (2018b; MathWorks, US), R (3.6.3; R Core Team, 2013), JAMOVI (1.1.9.0; The Jamovi Project, 2020), and JASP (0.12; JASP Team, 2018). Two people with CRPS were unable to complete the procedures with their affected hand. One of these participants (UL01) completed the full protocol with their non-affected hand. The other participant (UL15), however, was not able to do so due to the pain in her affected hand, and stopped after completing the third Open-loop Block in the washout phase (W-OL3) with her non-affected hand. As described in section 2.2.2, four participants with CRPS (UL01, UL03, UL11, UL13) completed additional trials in the washout phase. Because the number of washout trials can influence the measured retention of prism adaptation after-effects (Fernández-Ruiz & Díaz, 1999), we excluded the retention phase data of these four participants.

We used linear mixed models regression for our main analyses, because this method allowed us to include all available data from all participants regardless of the missing elements just described. This analysis also benefits from being robust against violations of distributional assumptions (Schielzeth et al., 2020). We also used mixed ANOVAs for supplementary analyses. To be concise, we only report interactions from analyses that address our hypotheses (i.e. that involve Group). We follow-up any significant three-way interactions to enable Group by Hand comparisons (e.g. within each level of Phase). We computed *p*-values from the linear mixed models regression with the Satterthwaite degrees of freedom (*lme4*, and *lmerTest* R packages; Bates, Mächler, Bolker, & Walker, 2015; Kuznetsova, Brockhoff, & Christensen, 2017), which has a lower Type-1 error rate than other methods (Luke, 2017). We report the ANOVA outputs from the linear mixed models analyses for ease of interpretation. We followed up any significant interactions with pairwise comparisons of the estimated marginal means derived from the linear mixed model (*emmeans*, and *multcomp* R packages; Hothorn, Bretz, & Westfall, 2008; Lenth, Singmann, Love, Buerkner, & Herve, 2019), also using the Satterthwaite degrees of freedom. For these, and pairwise comparisons stemming from the ANOVAs, we applied Holm-Bonferroni corrections (Holm, 1979) for multiple comparisons, indicated by “*p*_adjusted_”. We considered *p*-values < .05 as statistically significant. Calculating effect sizes for linear mixed models is complicated by the challenges associated with estimating the degrees of freedom, and partitioning variance in linear mixed models (Rights & Sterba, 2019). To overcome issues associated with points estimates of effect sizes, 95% confidence intervals are presented alongside effect size estimates (Pek & Flora, 2018). For the follow-up tests of linear mixed models where we performed pairwise comparisons, we calculated Cohen’s *d* from the estimated marginal means, the population standard deviation estimated by the linear mixed model, and the Satterthwaite degrees of freedom (*emmeans* R package). For results that are reported in short, and as part of a cluster (e.g. *t*s(51) ≤ 2.42, *p*s_adjusted_ ≥ .340), the effect size estimate is reported using absolute values.

See preregistration for a full list of planned analyses (https://osf.io/6jpfg/). Analyses that were not preregistered are specified as exploratory.

## 3. Results

### 3.1 Summary statistics

Descriptive statistics for clinical data and questionnaire measures for people with CRPS are presented in Table 2. As prism adaptation can be influenced by the speed of movement (Redding et al., 2005), and its main outcome measures relate to the precision of pointing movements, we conducted two-way mixed ANOVAs on peak velocity, peak acceleration, and baseline closed-loop endpoint errors, with Group and Hand as factors. These revealed that there was a tendency for peak velocity and peak acceleration to be smaller for people with CPRS when using their affected hand relative to their non-affected hand, although only the differences in peak acceleration were significant after correcting for multiple comparisons. There were no differences that depended on Group or Hand, or any interactions for baseline closed-loop endpoint errors (Supplementary Text T1).

In total, 17 people with CRPS and 18 controls completed the prism adaptation protocol. Two of the people with CRPS were unable to use their CRPS-affected hand and completed the experiment only with their non-affected hand. All other participants completed the paradigm with both hands (one hand at a time). Mean endpoint errors are presented in Fig. 4.

**Figure 4.**
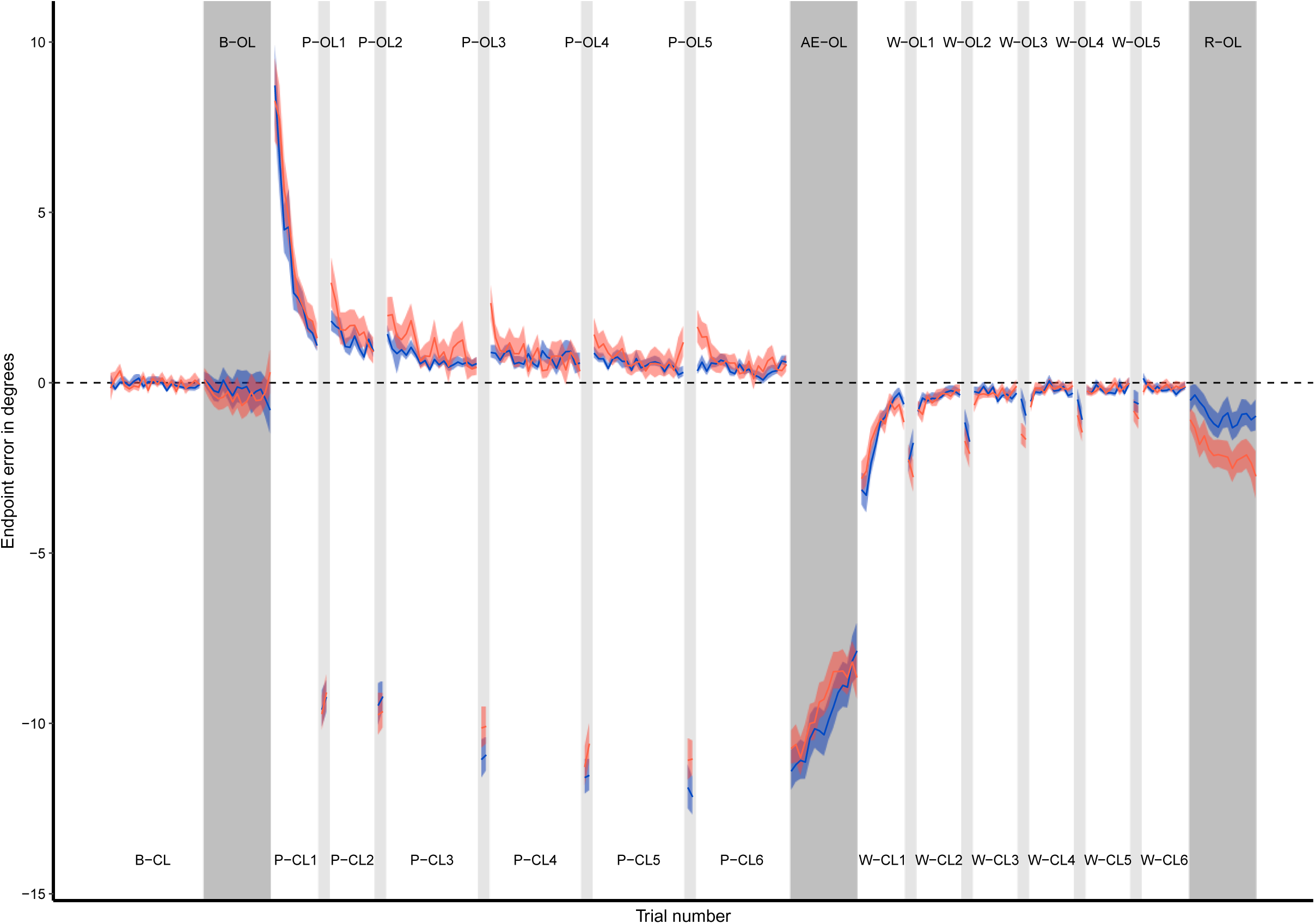
Endpoint errors in degrees are presented for people with CRPS (n = 17; red) and controls (n = 18; blue). Two people with CRPS only have data for their non-affected hand, and the data from four people with CRPS has been removed from the retention phase due their having completed additional trials during the washout phase (see main text for details; 2.2.2). The boundaries of the coloured shaded areas show the standard error of the mean. The grey shaded areas indicate open loop (OL) trials (i.e. when participants performed pointing movements without vision of their hand), for Open-loop Blocks (dark grey), Interim Open-loop Blocks (lighter grey), and Washout Open-loop Blocks (light grey). The white areas indicate closed-loop pointing, where participants had terminal exposure (i.e. they had vision of their hand toward the end of a movement). The black dashed line shows the target orientation (i.e. zero degree error). Negative values indicate endpoint errors made towards the affected/non-dominant side. AE = after-effect phase; B = baseline phase; CL = closed-loop pointing; OL = open-loop pointing; P = prism adaptation phase; R = retention phase; W = washout phase

### 3.2 Closed-loop endpoint errors

#### 3.2.1 Exponential decay of endpoint errors during prism exposure

To test whether people with CRPS would show impaired strategic control relative to controls (Hypothesis 1), and/or depend on the hand used (Hypothesis 5), we analysed the strategic error reduction during closed-loop trials during the prism exposure phase (i.e. P-CL1 – P-CL6) using linear mixed models regressions.

Before analysing the constants derived from the fitted models, we analysed the model fit parameters. The model failed to converge, or there was no exponential fit for one person with CRPS (non-affected hand), and for one control (dominant hand). When we analysed the model fit parameters (i.e. root-mean-square error, *adj. R^2^*) we found no significant effects of Group, or Hand, and no significant interactions (see Supplementary Text T2). Therefore, we proceeded to analyse the constants derived from the models (i.e. 1/*b*, and *c*; Fig. 5).

**Figure 5.**
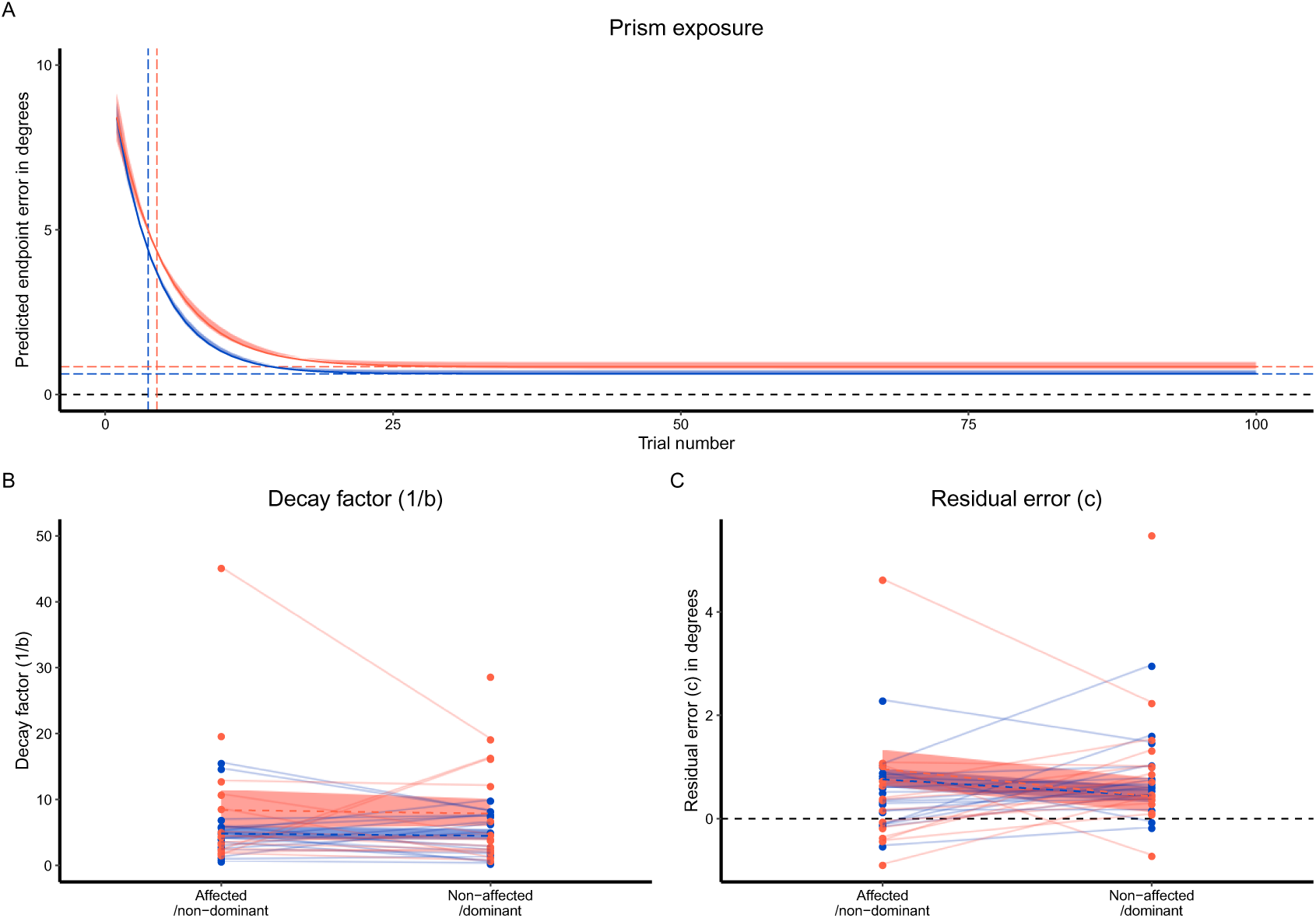
Exponential decay functions (*x* = *a* × *exp*^-*b* × *n*^ + *c*) fitted to averaged data (A) and individual decay factors (B) and residual errors (C) for end-point errors for the 100 closed-loop trials during the prism exposure phase. The decay factor (i.e. 1/*b*) shows the rate at which a fitted curve decays, and thus a large value indicates that more trials are needed for endpoint errors to reach the asymptote (i.e. the residual error, *c*). In panel A, solid lines indicate the predicted value, and the boundaries of the shaded areas depict the 95% confidence interval for the constants (i.e. *a*, *b*, *c*) fitted to group-level data (i.e. mean endpoint errors for each trial, split by Group). The black dashed lines (A, C) show perfect performance (i.e. zero degree error). Negative values indicate endpoint errors made towards the affected/non-dominant side (A, C). The coloured dashed lines (A) indicate the decay constant (i.e. 1/*b*; vertical lines), and the residual error (i.e. *c;* horizontal lines), for people with CPRS (red), and controls (blue). Points depict individual level data is presented for the decay factor (i.e. 1/*b*; B) and the residual error (i.e. *c*; C) for people with CRPS (n_affected_ = 15; n_non-affected_ = 16; red), and controls (n_non-dominant_ = 18; n_dominant_ = 17; blue). Data points are connected for each participant, given that they had data available and that we were able to fit it to an exponential decay function. The coloured dashed lines (B, C) indicate group means, and the boundaries of the coloured shaded areas show ± one Standard Error of the Mean.

The rate at which endpoint errors decayed did not differ between people with CRPS and controls, or between the affected/non-dominant and non-affected/dominant hand (Table 3). That is, there was no significant difference between people with CRPS (*M*_1/*b*_ = 8.64, *95% CI* [5.39, 11.89]) and controls (*M*_1/*b*_ = 4.62, *95% CI* [1.50, 7.74]) on the decay factor (i.e. 1/*b*). There was also no significant main effect of Hand (affected/non-dominant *M*_1/*b*_ = 6.94, *95% CI* [4.38, 9.50]; non-affected/dominant *M*_1/*b*_ = 6.32, *95% CI* [3.77, 8.87]). Furthermore, there was no significant interaction between Group and Hand. These results therefore indicate that there was no evidence of a difference in the rate of endpoint error decay during prism exposure for people with CPRS and controls, while using either hand.

**Table 3.** Inferential statistics for the analysis of decay factors (i.e. 1/b) and residual error (i.e. c) from closed-loop pointing during the prism exposure phase, examining the effect of Group, and Hand.

| Analysis | Result |  |
| --- | --- | --- |
| Decay factor |  |  |
| Group | $F(1, 30.29) = 3.33, p = .078$ | |
| Hand | $F(1, 28.19) = 0.33, p = .629$ | |
| Group x Hand | $F(1, 28.19) = 0.03, p = .860$ | |
| Residual error |  |  |
| Group | $F(1, 26.29) = 0.19, p = .669$ | |
| Hand | $F(1, 24.06) = 4.04, p = .056$ | # |
| Group x Hand | $F(1, 24.05) < 0.01, p = .949$ | |

The residual endpoint error during prism exposure was not found to differ between people with CRPS and controls (Table 3). That is, there was no significant difference in the residual error between people with CRPS (*M_c_* = 0.75, *95% CI* [0.26, 1.25]) and controls (*M_c_* = 0.61, *95% CI* [0.14, 1.08]). There was a tendency towards greater residual errors in the direction of the prismatic shift (i.e. towards the non-affected/dominant side) for the non-affected/dominant arm (*M_c_* = 0.86, *95% CI* [0.48, 1.24]) compared to the affected/non-dominant arm (*M_c_* = 0.50, *95% CI* [0.12, 0.88]), although not significant. There was no significant interaction between Group and Hand on the residual error. Our results therefore suggest that the residual error that remained once pointing correction had reached asymptote during prism exposure trials was not found to differ between people with CRPS and controls, and that there were no differences between groups that depended on the hand that was used.

#### 3.2.2 Exponential decay of endpoint errors during washout

To further test whether people with CRPS would show impaired strategic control relative to pain-free controls (Hypothesis 1), and/or depend on the hand used (Hypothesis 5), we analysed the strategic error reduction during closed-loop trials during the washout phase (i.e. W-CL1 – W-CL6) using linear mixed models regressions.

Prior to analysing the constants from the exponential decay function, we analysed the model fit. We were unable to fit an exponential decay function for two participants with CRPS, both for their affected hand. There were no difference in the model fits between people with CRPS and controls (i.e. no significant main effects or interactions involving Group for RMSE, *adj. R^2^*; see Supplementary Text T2), therefore we proceeded to analyse the constants derived from the models (i.e. 1/*b*, and *c*; Fig. 6).

**Figure 6.**
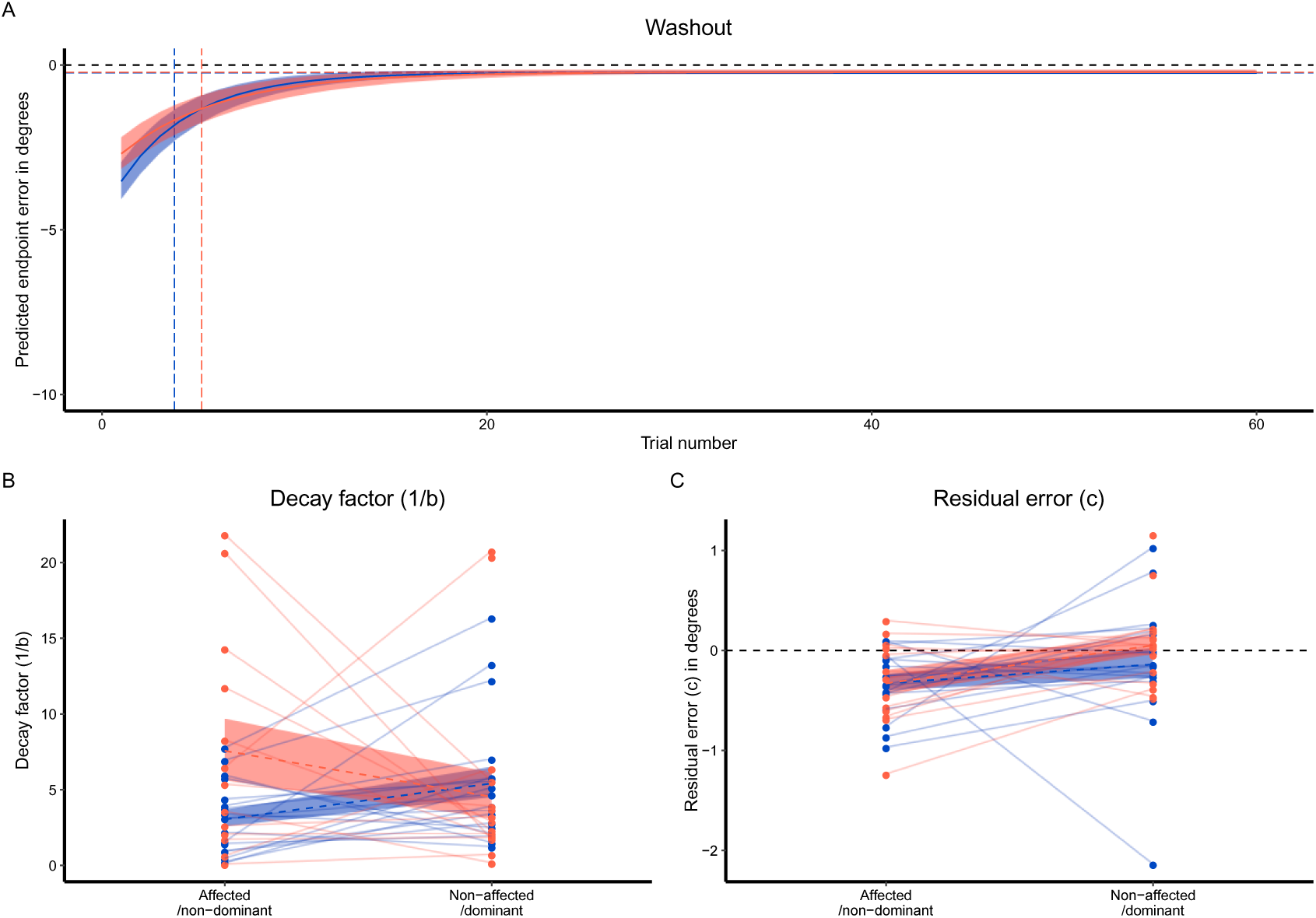
Exponential decay functions (*x* = *a* × *exp*^-*b* × *n*^ + *c*) fitted to averaged data (A) and individual decay factors (B) and residual errors (C) for end-point errors for the 60 closed-loop trials during the washout phase. The decay factor (i.e. 1/*b*) shows the rate at which a fitted curve decays, and thus a large value indicates that more trials are needed for endpoint errors to reach the asymptote (i.e. the residual error, *c*). In panel A, solid lines indicate the predicted value, and the boundaries of the shaded areas depict the 95% confidence interval for the constants (i.e. *a*, *b*, *c*) fitted to mean endpoint errors for each trial, split by Group. The black dashed lines (A, C) show perfect performance (i.e. zero degree error). Negative values indicate endpoint errors made towards the affected/non-dominant side (A, C). The coloured dashed lines (A) indicate the mean decay factor (i.e. 1/*b*; vertical lines), and the mean residual error (i.e. *c;* horizontal lines), for people with CPRS (red), and controls (blue). Points depict individual level data presented for the decay factor (i.e. 1/*b*; B) and the residual error (i.e. *c*; C) for people with CRPS (n_affected_ = 13; n_non-affected_ = 17; red), and controls (n_non-dominant_ = 18; n_dominant_ = 18; blue). Data points are connected for each participant, given that they had data available and that we were able to fit it to an exponential decay function. The coloured dashed lines (B, C) indicate group means, and the boundaries of the coloured shaded areas show ± one Standard Error of the Mean.

The rate at which closed-loop endpoint errors reduced during the washout phase was not found to differ between people with CRPS and controls, or for the affected/non-dominant and non-affected/dominant hand (Table 4). That is, the decay rate did not significantly differ between people with CRPS (*M*_1/*b*_ = 6.09, *95% CI* [4.03, 8.15]) and controls (*M*_1/*b*_ = 4.23, *95% CI* [2.34, 6.12]). There was also no significant difference in decay rate that depended on the Hand (affected/non-dominant *M*_1/*b*_ = 5.34, *95% CI* [3.44, 7.23]; non-affected/dominant *M*_1/*b*_ = 4.98, *95% CI* [3.22, 6.74]).

**Table 4.** Inferential statistics for the analysis of decay factors (i.e. 1/b) and residual error (i.e. c) from closed-loop pointing during the washout phase, examining the effect of Group, and Hand.

| Analysis | Result |
| --- | --- |
| Decay factor |  |
| Group | $F(1, 28.69) = 1.85, p = .184$ |
| Hand | $F(1, 28.12) = 0.09, p = .773$ |
| Group x Hand | $F(1, 28.12) = 5.04, p = .033$ * |
| Residual error |  |
| Group | $F(1, 62) = 0.62, p = .433$ |
| Hand | $F(1, 62) = 6.09, p = .016$ * |
| Group x Hand | $F(1, 62) = 0.63, p = .433$ |

\* $p < .05$ .

There was a significant interaction between Group and Hand on the decay rate, (Table 4; Fig. 5B). Although none of the follow-up comparisons were significant, the interaction appeared to be driven by a tendency for people with CRPS to have a larger decay factor when using the affected hand (*M*_1/*b*_ = 7.63, *95% CI* [4.75, 10.52]) than controls when using the non-dominant hand (*M*_1/*b*_ = 3.04, *95% CI* [0.58, 5.52]), *t*(51) = 2.42, *p*_adjusted_ = .340, *d* = 0.94*, 95% CI* [0.14, 1.74]. The decay rates were numerically more similar, and the difference was in the opposite direction, for the non-affected/dominant hand (CRPS *M*_1/*b*_ = 4.54, *95% CI* [2.02, 7.07]; controls *M*_1/*b*_ = 5.42, *95% CI* [2.96, 7.87]), *t*(51) = 0.50, *p*_adjusted_ = .921, *d* = −0.18*, 95% CI* [−0.90, 0.55]. This result indicates that there was a non-significant tendency for people with CRPS to need more trials to reduce their endpoint errors during washout when using their affected hand than controls when using their non-dominant hand, which might indicate that coordinating motor control takes longer for the CRPS affected-hand.

The residual endpoint error during washout trials did not differ between people with CRPS and controls (Table 4). That is, there was no main effect of Group (CRPS *M_c_* = −0.14, *95% CI* [−0.32, 0.33]; controls *M_c_* = −0.24, *95% CI* [−0.39, −0.08]) on residual errors (i.e. *c*). In contrast, the residual error was different between hands. Participants had a greater magnitude of residual error towards the affected/non-dominant side (i.e. the direction opposite to the prismatic shift) for their affected/non-dominant hand (*M_c_* = −0.33, *95% CI* [−0.51, −0.16]) compared to their non-affected/dominant hand (*M_c_* = −0.04, *95% CI* [−0.20, 0.12]). The interaction between Group and Hand on residual error was not significant. Therefore, our results provide no evidence of a difference in residual error during washout between people with CRPS and controls. Participants had greater residual error in the direction opposite to the prismatic shift for their affected/non-dominant hand, compared to their non-affected/dominant hand, which did not vary between Groups.

### 3.3 Open-loop endpoint errors

#### 3.3.1 Prism adaptation after-effects

To address the hypothesis that people with CRPS would show impaired sensorimotor adaptation relative to controls (Hypothesis 2), and/or that it would differ depending on the hand used (Hypothesis 5), we used a linear mixed models regression to analyse the effect of Group, Hand, and Phase (baseline, after-effect, retention) on open-loop pointing endpoints (analysis PAE, Fig. 2, Fig. 4).

Participants adapted to the prismatic shift introduced by the goggles and showed some retention of this effect after the washout phase. There was a significant main effect of Phase, *F*(2, 42.93) = 1077.62, *p* < .001. Errors in the open loop trials performed during the after-effect phase were significantly deviated in the direction opposite to the prismatic shift (*M* = −9.77°, *95% CI* [−10.36, −9.17]) compared to both the baseline phase (*M* = −0.34°, *95% CI* [−0.93, 0.26]), *t*(43) = 42.61, *p*_adjusted_ ≤ .001, *d* = 4.07*, 95% CI* [3.16, 4.96], and the retention phase (*M* = - 1.46°, *95% CI* [−2.06, −0.86]), *t*(43) = 37.04, *p*_adjusted_ ≤ .001, *d* = −3.58*, 95% CI* [−4.38, −2.78]. Endpoint errors were also significantly deviated for the retention phase compared to the baseline phase, *t*(43) = 5.01, *p*_adjusted_ < .001, *d* = 0.48*, 95% CI* [0.26, 0.71]. These results indicate that adaptation to prismatic visual shifts produced sensorimotor after-effects (i.e. open-loop endpoint errors biased in the direction opposite to the prismatic shift relative to baseline), and that there was some retention of this effect after participants completed washout trials.

There was also a significant main effect of Hand on the open-loop endpoint errors, *F*(1, 2768.12) = 114.97, *p* < .001, whereby participants made greater errors towards their affected/non-dominant side (i.e. in the direction opposite to the prismatic shift) when using their affected/non-dominant hand (*M* = −4.33°, *95% CI* [−4.88, −3.78]) than with their non-affected/dominant hand (*M* = −3.38°, *95% CI* [−3.93, −2.83]). However, there was no main effect of Group (CRPS *M* = −4.00°, *95% CI* [−4.77, −3.23]; controls *M* = −3.71°, *95% CI* [−4.45, −2.96]), *F*(1, 32.63) = 0.32, *p* = .576.

There was a significant interaction between Group and Phase, *F*(2, 2766.60) = 21.21, *p* < .001. This interaction was superseded by a significant interaction between Group, Hand, and Phase, *F*(2, 2762.70) = 22.37, *p* < .001 (Table 5).

**Table 5.** Estimated marginal means and corresponding confidence intervals for endpoint errors made during open-loop trials, split by Group, Hand, and Phase (baseline, prism adaptation after-effects, retention).

| | Affected/non-dominant | | Non-affected/dominant | | $p_{\text{Hand}}$ |
| --- | --- | --- | --- | --- | --- |
| | $M$ | 95% CI | $M$ | 95% CI | |
| Baseline |  |  |  |  |  |
| CRPS | -0.28° | [-1.12, 0.57] | -0.64° | [-1.47, 0.20] |  |
| Controls | -0.73° | [-1.54, 0.08] | 0.30° | [-0.52, 1.11] | *** |
| $p_{\text{Group}}$ | | | | | |
| After-effect |  |  |  |  |  |
| CRPS | -10.43° | [-11.27, -9.59] | -8.70° | [-9.53, -7.86] | *** |
| Controls | -10.17° | [-10.98, -9.35] | -9.77° | [-10.59, -8.96] |  |
| $p_{\text{Group}}$ | | | | | |
| Retention |  |  |  |  |  |
| CRPS | -2.54° | [-3.40, -1.68] | -1.42° | [-2.28, -0.55] | *** |
| Controls | -1.83° | [-2.65, -1.02] | -0.05° | [-0.86, 0.78] | *** |
| $p_{\text{Group}}$ | # | | | | |

$p_{\text{Group}}$ = adjusted $p$ -values for between-group comparisons; $p_{\text{Hand}}$ = adjusted $p$ -values for within-group comparisons of the Hand used; \*\*\* $p < .001$ ; # $p = .051$ .

We followed up this interaction by performing four pairwise comparisons between each level of Group and Hand per Phase (baseline, after-effects, retention; Fig. 7). For the baseline Phase, control participants made greater errors towards their non-dominant side with their non-dominant hand compared to their dominant hand, *t*(950) = 5.04, *p*_adjusted_ < .001, *d* = 0.44*, 95% CI* [0.27, 0.62]. There was no such difference for people with CRPS, and no significant differences between Groups for either Hand, *ts*(950) ≤ 1.70, *p*s_adjusted_ ≥ .663, *d*s ≤ 0.40. During the after-effect Phase, people with CRPS deviated further towards their affected side (i.e. the direction opposite to the prismatic shift) with their affected hand than their non-affected hand, *t*(950) = 7.97, *p*_adjusted_ < .001, *d* = 0.75*, 95% CI* [0.55, 0.94]. There was no significant difference between Hands for controls participants, and no differences between Groups for either hand, *ts*(950) ≤ 1.96, *p*s_adjusted_ ≥ .381, *d*s ≤ 0.46. In the retention phase, control participants made greater errors with their non-dominant hand than their dominant hand, *t*(950) = 8.35, *p*_adjusted_ < .001, *d* = 0.77, *95% CI* [0.59, 0.95]). Similarly, people with CRPS made greater errors with their affected hand than their non-affected, *t*(950) = 4.55, *p*_adjusted_ < .001, *d* = 0.49, *95% CI* [0.27, 0.70]. Comparing Groups, there was a tendency for people with CRPS to make greater errors in the direction opposite to the prismatic shift when using their non-affected hand than controls using their dominant hand. This difference was not significant after controlling for multiple comparisons *(t*(950) = 2.45, *p*_adjusted_ = .051), however the effect size was large *(d* = 0.59, *95% CI* [0.11, 1.07]). There was no significant difference between Groups when using their affected/non-dominant hand (*t*(950) = 1.28, *p*_adjusted_ = .263), with a medium effect size (*d* = 0.31, *95% CI* [−0.18, 0.79]). These results suggest that each Group had a tendency for larger (i.e., more negative) endpoint errors for their affected/non-dominant hand than their non-affected/dominant hand in at least two phases, and that there was a weaker evidence for this tendency for people with CRPS (compared to controls) to have larger endpoint errors persisting through the retention phase when using their non-affected/dominant hand.

**Figure 7.**
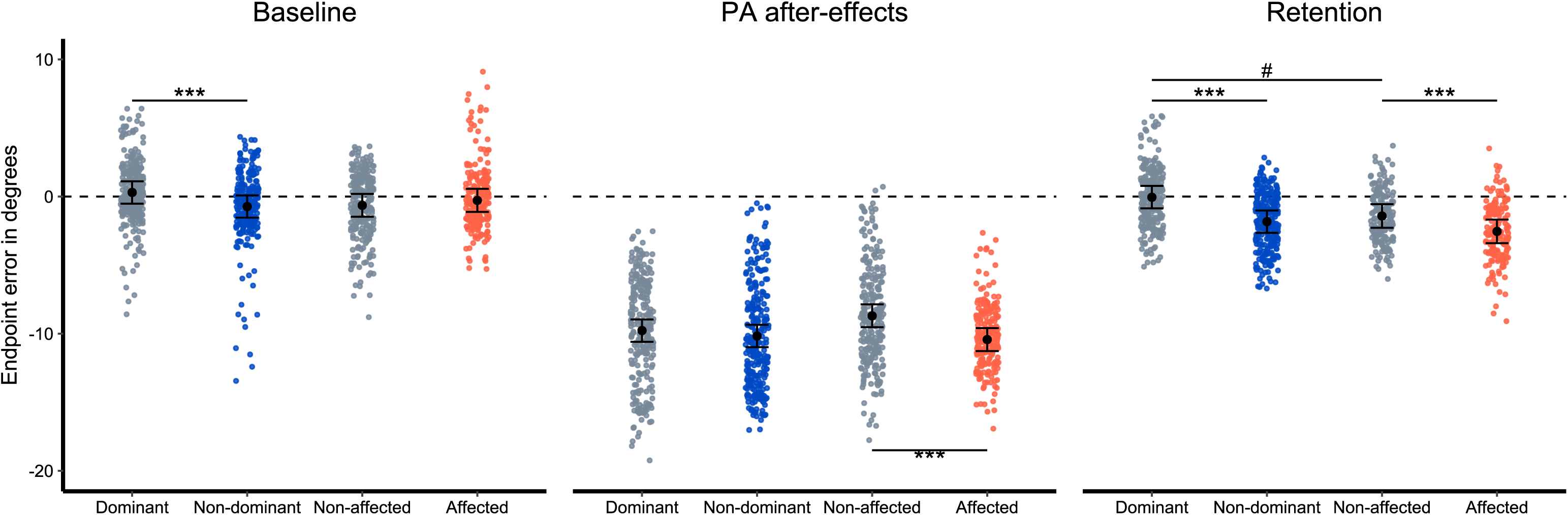
Open-loop endpoint errors for the baseline (B-OL), after-effect (AE-OL), and retention (R-OL) phases are presented for people with CRPS (n = 17; red and grey) and controls (n = 18; blue and grey), for the affected (red)/non-dominant (blue) and the non-affected/dominant (grey) hands. Two people with CRPS only have data for their non-affected hand, and the data from four people with CRPS has been removed from the retention phase due to having completed an additional 42 washout trials. Endpoint errors (in degrees) are presented for individual trials included in the linear mixed models regression analysis (i.e. for the 15 open-loop trials for each Open-loop Block, for each Hand, and for each participant). Black points and error bars depict the estimated marginal means and 95% confidence intervals derived from the analysis. The black dashed line shows the target orientation (i.e. zero degree error). Negative values indicate endpoint errors made towards the affected/non-dominant side. PA = prism adaptation. # *p*_adjusted_ = .051, *** *p*_adjusted_ < .001

To confirm that these results were not influenced by baseline open-loop pointing performance (i.e. “Absolute Baseline Error”; Supplementary Text T3), the direction of the prismatic shift, and/or the counterbalancing order, we ran three exploratory ANCOVAs with baseline open-loop pointing as a covariate, comparing between the direction of prismatic shift and/or the counter balancing order. We found no evidence to suggest that our results were influenced by any differences between participants in baseline open-loop pointing performance, *F*s(2, 62) ≤ 1.03, *p*s ≥ .362, ƞ^2^_p_ ≤ .03, the direction of the prismatic shift, *F*s(2, 62) ≤ 0.62, *p*s ≥ .544, ƞ^2^_p_ ≤ .02, or the counterbalancing order *Fs*(2, 62) ≤ 2.64, *p*s ≥ .080, ƞ^2^_p_ ≤ .08 (Supplementary Text T3). Furthermore, visual exploration of the data provided no indication that our results were influenced by previous experience with prism adaptation (Figs. S1 & S2).

#### 3.3.2 Development of prism adaptation after-effects

To address the hypothesis that sensorimotor realignment might develop at a different rate for people with CRPS, compared to controls (Hypothesis 3), and/or depend on the hand used (Hypothesis 5), we used a linear mixed models regression to analyse the effects of Group and Hand on endpoint errors during the Open-loop Blocks of the prism exposure phase (Fig. 2 and Fig. 4; P-OL1 – P-OL5).

There was a main effect of Open-loop Block on endpoint errors, *F*(4, 617.25) = 23.50, *p* < .001. We followed this effect up by comparing each Interim Open-loop Block (P-OL2 to P-OL5) to the first block (i.e. P-OL1; see Table 6). This effect was driven by a greater magnitude of endpoint errors made during the fourth and fifth blocks compared to the first block. The second and third blocks were not significantly different to the first block.

**Table 6.**
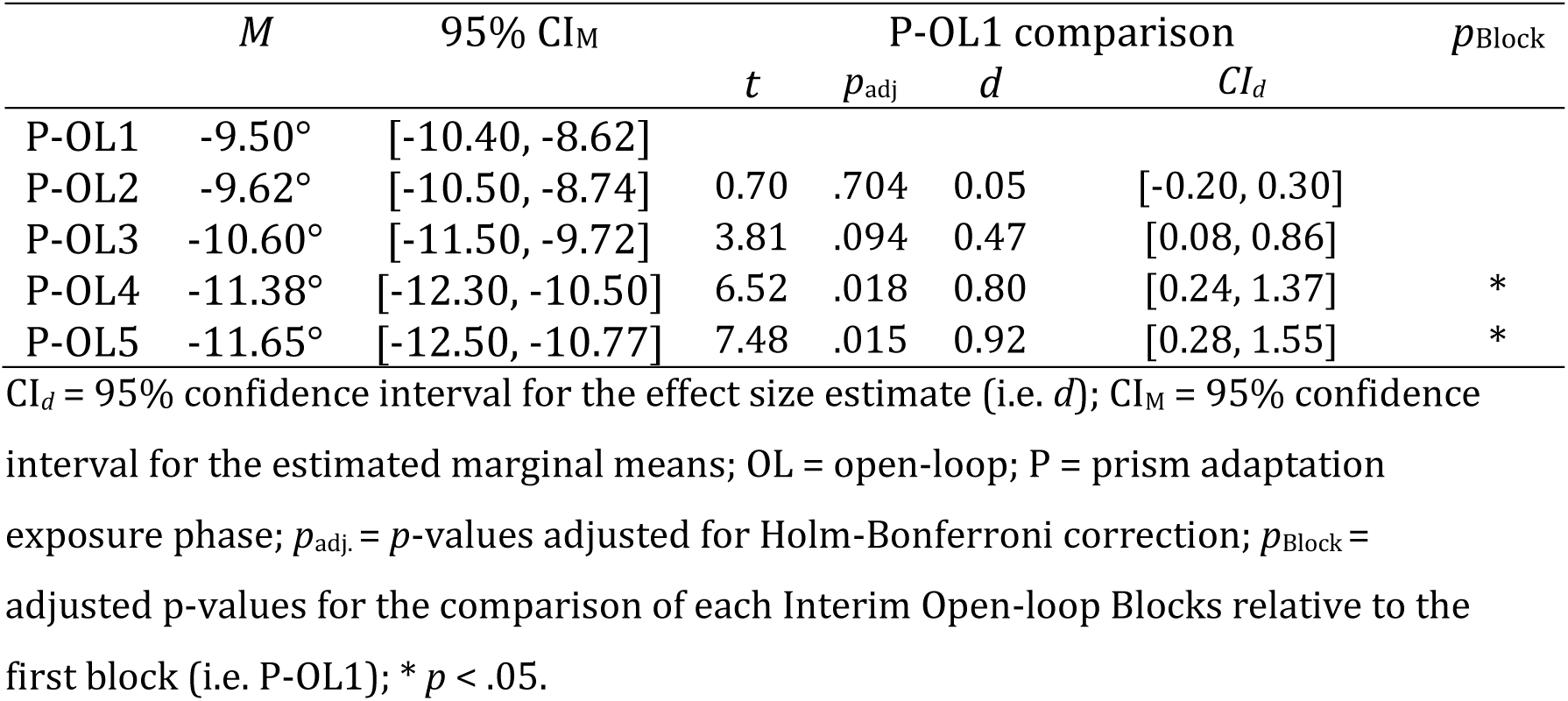
Descriptive and inferential statistics presented for endpoint errors made on Open-Loop Blocks during the prism exposure phase (ΔP-OL).

There was also a significant main effect of Hand, *F*(1, 620.79) = 8.32, *p* = .003, whereby participants made greater errors towards the affected/non-dominant side (i.e. in the direction opposite to the prismatic shift) when using their affected/non-dominant hand (*M* = −10.80°, *95% CI* [−11.70, −9.99]) compared to their non-affected/dominant hand (*M* = −10.30°, *95% CI* [−11.10, −9.45]). There was no significant main effect of Group, and no significant interactions *F*s(4, 620.79) ≤ 2.27, *p* ≥ .060. These findings provide no evidence of a difference for development of prism-adaptation after-effects between people with CRPS and controls, and that this did not vary depending on the hand being used.

#### 3.3.3 Decline of prism adaptation after-effects

To address the hypothesis that sensorimotor realignment might decline at a different rate for people with CRPS, compared to controls (Hypothesis 3), and/or depend on the hand used (Hypothesis 5), we used a linear mixed models regression to analyse the effect of Group and Hand on endpoint errors during the Open-loop Blocks of the washout phase (Fig. 4; W-OL1 – W-OL5).

The after-effects decayed during washout trials. This was evidenced by a main effect of Washout Open-loop Blocks on endpoint errors, *F*(4, 5.04) = 10.92, *p* = .011. We followed this effect up by comparing each Washout Open-loop Block (W-OL2 to W-OL5) to the first block (i.e. W-OL1; see Table 7). This effect was driven by a smaller magnitude of endpoint errors made during the fifth block than the first block. The second, third, and fourth blocks were not significantly different to the first block after correcting for multiple comparisons.

**Table 7.** Descriptive and inferential statistics presented for endpoint errors made on Open-Loop Blocks during the washout phase (ΔW-OL).

| | $M$ | 95% $CI_M$ | W-OL1 comparison | | | | $p_{Block}$ |
| --- | --- | --- | --- | --- | --- | --- | --- |
| | | | $t$ | $p_{adj}$ | $d$ | $CI_d$ | |
| W-OL1 | -2.30° | [-2.88, -1.72] |  |  |  |  |  |
| W-OL2 | -1.70° | [-2.27, -1.12] | 2.28 | .217 | -0.37 | [-0.79, 0.05] |  |
| W-OL3 | -1.20° | [-1.77, -0.62] | 5.84 | .077 | -0.68 | [-1.24, -0.11] |  |
| W-OL4 | -0.98° | [-1.56, -0.41] | 4.73 | .061 | -0.80 | [-1.43, -0.17] |  |
| W-OL5 | -0.76° | [-1.33, -0.18] | 4.97 | .044 | -0.94 | [-1.65, -0.23] | * |

$CI_d$ = 95% confidence interval for the effect size estimate (i.e. $d$ ); $CI_M$ = 95% confidence interval for the estimated marginal means; OL = open-loop; $p_{adj}$ = $p$ -values adjusted for Holm-Bonferroni correction; $p_{Block}$ = adjusted $p$ -values for the comparison of each Interim Open-loop Blocks relative to the first block (i.e. W-OL1); W = washout phase; \* $p < .05$ .

Participants had a greater retention of prism adaptation after-effects for their affected/non-dominant hand. That is, there was a significant main effect of Hand on endpoint errors during Washout Open-loop Blocks, *F*(1, 618.86) = 93.92, *p* < .001. Participants made greater errors in the direction opposite to the prismatic shift (i.e. towards the affected/non-dominant side) when using their affected/non-dominant hand (*M* = −2.01°, *95% CI* [−2.47, −1.54]) compared to their non-affected/dominant hand (*M* = −0.77°, *95% CI* [−1.23, −0.30]). There was no significant main effect of Group, and no significant interactions that involved Group, Hand, and/or Washout Open-loop Block, *F*s(4, 618.86) ≤ 1.63, *p*s ≥ .211. These findings suggest that the decay of prism adaptation after-effects did not differ between people with CRPS and controls, and did not vary depending on the Hand being used.

### 3.4 Kinematic changes during prism exposure

To address the hypothesis that people with CRPS would show less evidence of trial-by-trial changes in movement plans to compensate for the prismatic shift than controls (Hypothesis 4), we analysed the change in initial trajectory orientations during early trials (i.e. closed loop trials 1 to 10 during prism exposure; P-CL1) for each Group, and Hand.

Prior to analysing the detrended data for initial trajectory orientations, we inspected the model fit. We were unable to fit initial trajectory orientations to the exponential decay function for one person with CRPS (non-affected hand), and two controls (one non-dominant hand, one dominant hand). When we compared the model fit parameters for the remaining participants, the amount of variance explained by the models did not differ between people with CRPS and controls, or depending on the Hand used (Supplementary Text T2).

There was no evidence for trial-by-trial feedforward control for controls or people with CRPS, from either the analysis of *t*-values or *p*-values. That is, we did not find any evidence of a significant linear relationship between endpoint errors and changes in initial trajectory orientation for either Group using either Hand, *t*s(14) ≤ 1.43, *p*s_adjusted_ ≥ .174, *d*s ≤ 0.36. When we pooled the data for each Group, and explored it for each hand, we found evidence that *t*-values were different from zero for the non-affected/dominant hand (Supplementary Text T4). The analysis of *p*-values from individual correlations showed no evidence of a bias towards zero for either Hand (affected, non-affected, dominant, non-dominant), *D^+^* ≤ 0.18, *ps* ≥ .104. Therefore, only the data pooled across all participants provides evidence that trial-by-trial feedforward motor control was used to reduce endpoint errors for the non-affected/dominant hand during early prism exposure trials. However, there was no such evidence when data were considered separately for each Group from either *t*-values or from *p*-values.

## 4. Discussion

Our study was the first to characterise sensorimotor adaptation in people with CRPS. The results did not support our main hypotheses, as we found no evidence that prism adaptation is impaired for people with CRPS compared to controls when the two groups were directly compared. Instead, we found some indications that CRPS might lead to a *greater* propensity for sensorimotor realignment, because people with CRPS showed a larger magnitude of open-loop pointing errors for their affected hand than their non-affected hand after prism exposure, whereas there was no difference between hands for the controls. Generally, however, we found no differences between people with CRPS and controls on most prism adaptation measures, which we interpret as no consistent evidence of impaired sensorimotor adaptation in CRPS. Below we discuss the findings related to strategic control (4.1), sensorimotor realignment (4.2), and the retention of prism adaption after-effects (4.3). We also consider the theoretical (4.4.1) and clinical (4.4.2) implications of our findings, and potential limitations (4.5) of our study.

### 4.1 Strategic control

We found no evidence to suggest that strategic recalibration was disrupted for people with CRPS (Hypothesis 1). That is, we did not observe any difference between people with CRPS and controls on endpoint errors during prism exposure or during washout. We also did not find any group differences in the number of trials needed for endpoint errors to decay during prism exposure.

### 4.2 Sensorimotor realignment

Overall, we found no evidence that sensorimotor realignment was impaired for people with CRPS compared to controls (Hypotheses 2 & 3), and very little evidence of any group differences. We did not find any difference in the residual errors during prism exposure or during washout. There was no overall difference in the endpoint errors made during open-loop pointing directly after prism exposure. However, when the hands were considered separately, people with CRPS made greater endpoint errors in the prism adaptation after-effect phase with their affected hand than their non-affected hand, whereas there was no difference between hands for the controls in this phase. We interpret this finding tentatively, because direct comparisons between the groups revealed no differences in the open-loop pointing errors for the affected versus non-dominant and non-affected versus dominant hands. Nonetheless, these results suggest that, if anything, sensorimotor realignment was more pronounced for the affected hand, directly contradicting the idea that any impairment to sensorimotor adaptation in CRPS would be more/only evident when using this hand (Hypothesis 5).

More pronounced sensorimotor realignment in the affected hand could relate to lower stability of representations of the body and space. In a previous study, we showed that following tool use, people with upper limb CRPS showed more pronounced changes in their arm and peripersonal space representations compared to matched controls (Vittersø et al., 2020). Furthermore, in people with upper limb CRPS, representations of the affected and non-affected arms updated differently. Proprioceptive reference frames are closely linked to bodily representations (Medina & Coslett, 2010), and are updated during sensorimotor realignment (Jeannerod & Rossetti, 1993; Redding et al., 2005). Thus, the findings of the current study could be interpreted as further evidence that representations of the body and peripersonal space are less stable - or more malleable - in CRPS.

An outstanding issue is what mechanisms underly this malleability. One possibility is that it could relate to impaired online control due to the motor deficits that are diagnostic of CRPS. Differences in sensorimotor realignment between hands would be expected if participants used online control, and thus strategic control, to different extents for each hand when reducing endpoint error during prism exposure (Redding et al., 2005). The greater post prism adaptation open-loop pointing errors we observed for the affected hand (compared to the unaffected hand) suggests that participants may have relied less on strategic correction when using their affected hand, allowing for a greater proportion of error reduction to occur through sensorimotor realignment. We found, however, no evidence of any difference in strategic control between hands for the CRPS group, or between the participants with CRPS and the controls (4.1). Alternatively, the experience of pain could have influenced the motor learning involved in prism adaptation (Petitet, O’Reilly, & O’Shea, 2018) for people with CRPS. Using different paradigms to ours, such as a tracing or a repetitive typing task, past research has found greater motor learning during experimental pain induction relative to a placebo (Dancey, Murphy, Andrew, & Yielder, 2016a; Dancey, Murphy, Andrew, & Yielder, 2016b; Mavromatis, Neige, Gagné, Reilly, & Mercier, 2017), and for people with hand arthritis relative to pain-free controls (Parker, Lewis, Rice, & McNair, 2017). Experimental pain induction is thought to recruit additional attentional resources to the stimulated area and thereby facilitate motor learning (Dancey et al., 2016a; Dancey et al., 2016b). People with CRPS often report having to focus their attention on their affected limb in order to perform goal-directed movements, for instance on two of the questionnaires used in the present study (i.e. the Bath CRPS Body Perception Disturbance Scale, and the Neurobehavioral questionnaire; Galer & Jensen, 1999; Galer, Jensen, & Butler, 2013; Lewis & McCabe, 2010). Therefore, the increased malleability of sensorimotor representations that we observed for people with CRPS could relate to the pain that they experienced during the prism adaptation protocol, and the additional attentional demands that they were faced with.

### 4.3 Retention of prism adaptation after-effects

The decay of the prism adaptation after-effects was not found to differ between groups during open-loop washout trials. There was also no difference in the residual error during closed-loop washout trials between groups. There was a tendency for people with CRPS to retain prism adaptation after-effects for longer and/or to a greater extent than controls when using their non-affected/dominant hand, although this effect was not significant. Nonetheless, this pattern is worth noting because such a difference, if significant, would suggest that people with CPRS have more rigid sensorimotor representations of their non-affected hand. This is consistent with our finding that they showed a smaller magnitude of post prism adaptation open-loop errors for their non-affected hand than their affected hand. However, we did not find any evidence to suggest that the retention of prism adaptation after-effects was impaired for people with CRPS compared to controls.

### 4.4 Implications

#### 4.4.1 Theoretical implications

Harris’ (1999) sensorimotor theory of pain centres around the proposal that people with certain pathological pain conditions, such as CRPS (McCabe & Blake, 2007), experience sustained incongruence between sensory and motor information. Discrepancies between sensory and motor information that we experience in daily life are relatively minor, and can be resolved through sensorimotor adaptation (Wolpert et al., 2011). To experience sustained incongruence, people with CRPS and related conditions (e.g. fibromyalgia) should show less capacity to resolve such conflict through sensorimotor adaptation. However, we did not observe any impairment for people with CRPS in the magnitude of sensorimotor adaptation following exposure to a lateral visual distortion, nor in the rate at which the after-effect developed across exposure blocks. In fact, we observed some tentative indications that the painful limb has an enhanced propensity for sensorimotor realignment. Therefore, if pain is a result of conflicting sensorimotor information, then these findings provide no evidence that this conflict is driven by impaired sensorimotor adaptation for people with CRPS.

It should be noted, however, that our study focused on adaptation to sensorimotor conflict rather than the sensory and motor effects of such conflict when it is sustained (e.g. if adaptation is not possible). A recent study in people with fibromyalgia has suggested sensory abnormalities in chronic pain may arise due to an impaired ability to update forward models (i.e. the predicted sensory outcome of a movement), whilst inverse models (i.e. estimates of the current state of the body based on the available sensory information) are intact (Clementine Brun, McCabe, & Mercier, 2020). The sensorimotor realignment that occurs during prism adaptation is thought to cause both inverse and forward models to update (Petitet et al., 2018), however, our experiment was not designed to discriminate between these two types of changes. It is therefore possible that people with CRPS would have difficulties updating their forward models, which may not have been detected by our study due to the design and limited power. Our findings, however, do not suggest that updating of sensorimotor representations is impaired for people with CRPS. Intact sensorimotor updating should enable people with CRPS to compensate for incongruent sensory and motor information, and thus reduce the likelihood of the hypothesised sensorimotor incongruence to occur (Harris, 1999). Therefore, our findings contradict an implicit assumption of the sensorimotor theory of pain, or at least suggest that the sensorimotor conflict that this model is centred around is unlikely to be related to impaired adaptation.

#### 4.4.2 Clinical implications

Prism adaptation is considered a promising treatment for hemispatial neglect following brain damage (Luaute, Halligan, Rode, Jacquin-Courtois, & Boisson, 2006; Rossetti, Kitazawa, & Nijboer, 2019). This, along with early reports of deviations of subjective body midline towards the affected side (Sumitani et al., 2014; Sumitani, Rossetti, et al., 2007; Sumitani, Shibata, et al., 2007; Uematsu et al., 2009), and attention bias away from the affected limb in CRPS (Bultitude, Walker, & Spence, 2017; Filbrich et al., 2017; Moseley, Gallace, & Iannetti, 2012; Moseley, Gallace, & Spence, 2009; Reid et al., 2016), led to investigations of prism adaptation as a treatment for CRPS (Bultitude & Rafal, 2010; Christophe et al., 2016; Sumitani, Rossetti, et al., 2007). However, in the first double-blind randomized controlled trial of prism adaptation treatment for CRPS, we recently found no benefit of prism adaptation compared to sham treatment (Halicka, Vittersø, et al., 2020b). The results of the present study indicate that this lack of improvement cannot be attributed to problems adapting to the prismatic shift. This, as well as other recent findings that people with CRPS showed no spatial attention bias (e.g. De Paepe et al., 2020; Filbrich et al., 2017; Halicka, Vittersø, et al., 2020a; Ten Brink, Halicka, Vittersø, Keogh, & Bultitude, 2020), leads us to conclude that prism adaptation is ineffective because spatial representation and attention do not underlie the physical manifestations of CRPS. Nonetheless, our findings that sensorimotor adaptation is no different to – perhaps even better than – normal suggests that sensorimotor processing could be exploited for rehabilitation.

### 4.5 Limitations

Although comparable in size to past research into people with CRPS (e.g. Clémentine Brun, Giorgi, et al., 2019; De Paepe et al., 2020; Filbrich et al., 2017; Verfaille et al., 2020), our total sample size may have limited the power to detect effects, especially for analyses where we had to exclude datapoints, such as for the analysis of decay rates. Our sample of people with CRPS was heterogeneous, and included people with additional painful conditions, some of which can be bilateral. However, these are common comorbidities in CRPS (Beerthuizen et al., 2012). Furthermore, 8 of the people with CRPS had previous experience with prism adaptation (two weeks, twice daily, using the affected hand) as part of a double-blind randomized controlled trial (Halicka, Vittersø, et al., 2020b). Prior experience of prism adaptation can influence performance during a subsequent prism adaptation session, by reducing the magnitude of adaptation (Martin et al., 1996). A study of patients with hemispatial neglect following stroke that used a similar two-week prism adaptation regimen found significant sensorimotor after-effects at 12 hours post-treatment, but not at 84 hours post-treatment (Frassinetti, Angeli, Meneghello, Avanzi, & Làdavas, 2002). It is not known whether prism adaptation after-effects from repeated sessions carry over for longer than this in people without brain damage. However, we consider it unlikely that they would persist for as long as 7 months or more (the minimum time between studies for the participants who took part in both the randomized controlled trial and the present study). Our sample size did not permit us to make any direct comparisons between those who had previous experience with prism adaptation and those who did not, although visual exploration of the data did not suggest that our main findings were influenced by previous experience with prism adaptation (Figs. S1 & S2). Furthermore, there was no evidence of reduced sensorimotor adaptation in people with CRPS (regardless of prior experience with prism adaptation). For these reasons, it is unlikely that there were any additive effects of previous prism exposure, although we cannot rule out that participants remembered a strategy for compensating for the lateral optical shift. In this case, our current findings would underestimate the sensorimotor after-effects in CRPS. Therefore, our results would still not support the hypothesised impairment in sensorimotor realignment.

## 5. Conclusions

We found no evidence of impaired strategic recalibration or sensorimotor realignment in people with CRPS, though they showed greater prism adaptation after-effects for their affected hand than for their non-affected hand. These findings contradict our hypotheses, based on the sensorimotor theory of pain, that people with CRPS would have impaired adaptation and that any deficit would be more pronounced when using the affected limb. Instead, our findings add to previous evidence that people with CRPS show a more pronounced updating of bodily and peripersonal space representations, than pain-free individuals. Our results challenge an implicit assumption of existing theories of how pain might be maintained in the absence of clear tissue pathology, and add to our understanding of neuropsychological changes in CRPS.

## Acknowledgements

The authors thank Ms Eve Evans for her help with the study, Prof Andrea Serino, Dr Karin Petrini, two anonymous reviewers for their insightful comments on the manuscript, and the participants for volunteering their time.

## 6. Supplementary material

### 6.1 Supplementary Figure S1: Closed-loop pointing by previous prism exposure

**Figure S1.**
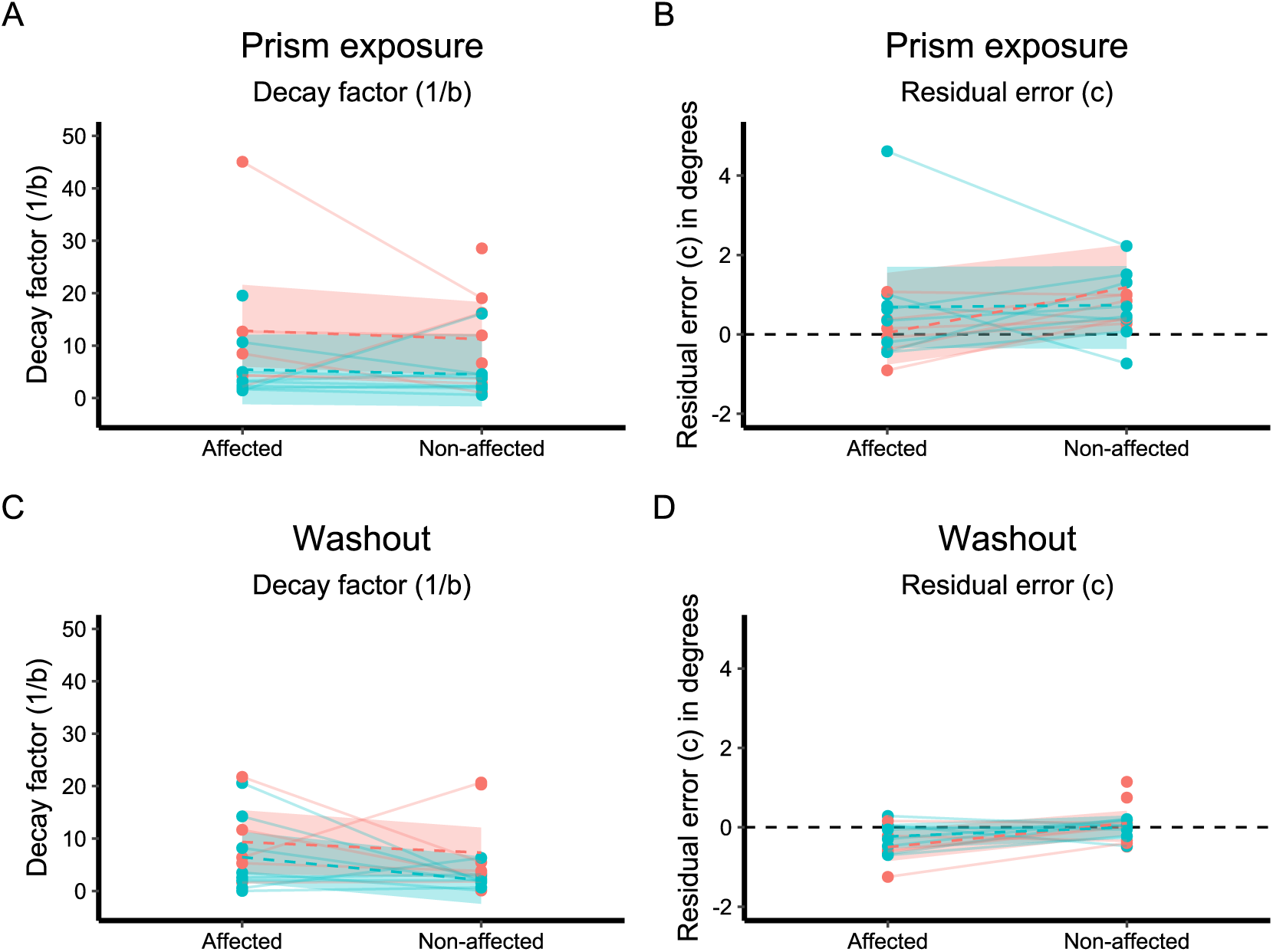
Individual decay factors (A, C) and residual errors (B, D) from closed-loop trials during the prism exposure phase (A, B) and the washout phase (C, D) are depicted, split by previous experience with prism adaptation. Eight of the participants with CRPS (red) had previous experience with prism adaptation from a two-week intervention (Halicka, Vittersø, et al., 2020b) ≥ 7 months prior to taking part in the present study. The remaining nine people with CRPS (blue) had no previous experience with prism adaptation, as they either received sham treatment or were not enrolled in the trial. The decay factor (i.e. 1/*b*) shows the rate at which a fitted curve decays, and thus a large value indicates that more trials are needed for endpoint errors to reach the asymptote (i.e. the residual error, *c*). The black dashed lines (B, D) show perfect performance (i.e. zero degree error). Negative values indicate endpoint errors made towards the affected/non-dominant side (B, D). The coloured dashed lines indicate estimated marginal means, and the boundaries of the coloured shaded areas show the upper and lower boundaries of 95% confidence intervals for the estimated marginal means. Points depict individual level data for the decay factor (i.e. 1/*b*; A, C) and the residual error (i.e. *c*; B, D). Data points are connected for each participant, given that they had data available and that we were able to fit it to an exponential decay function to data from the prism exposure phase (previous prism experience: n_affected_ = 6, n_non-affected_ = 8; no experience/sham: n_affected_ = 9, n_non-affected_ = 8) and the washout phase (previous prism experience: n_affected_ = 5, n_non-affected_ = 8; no experience/sham: n_affected_ = 8, n_non-affected_ = 9).

### 6.2 Supplementary Figure S2: Open-loop pointing by previous prism exposure

**Figure S2.**
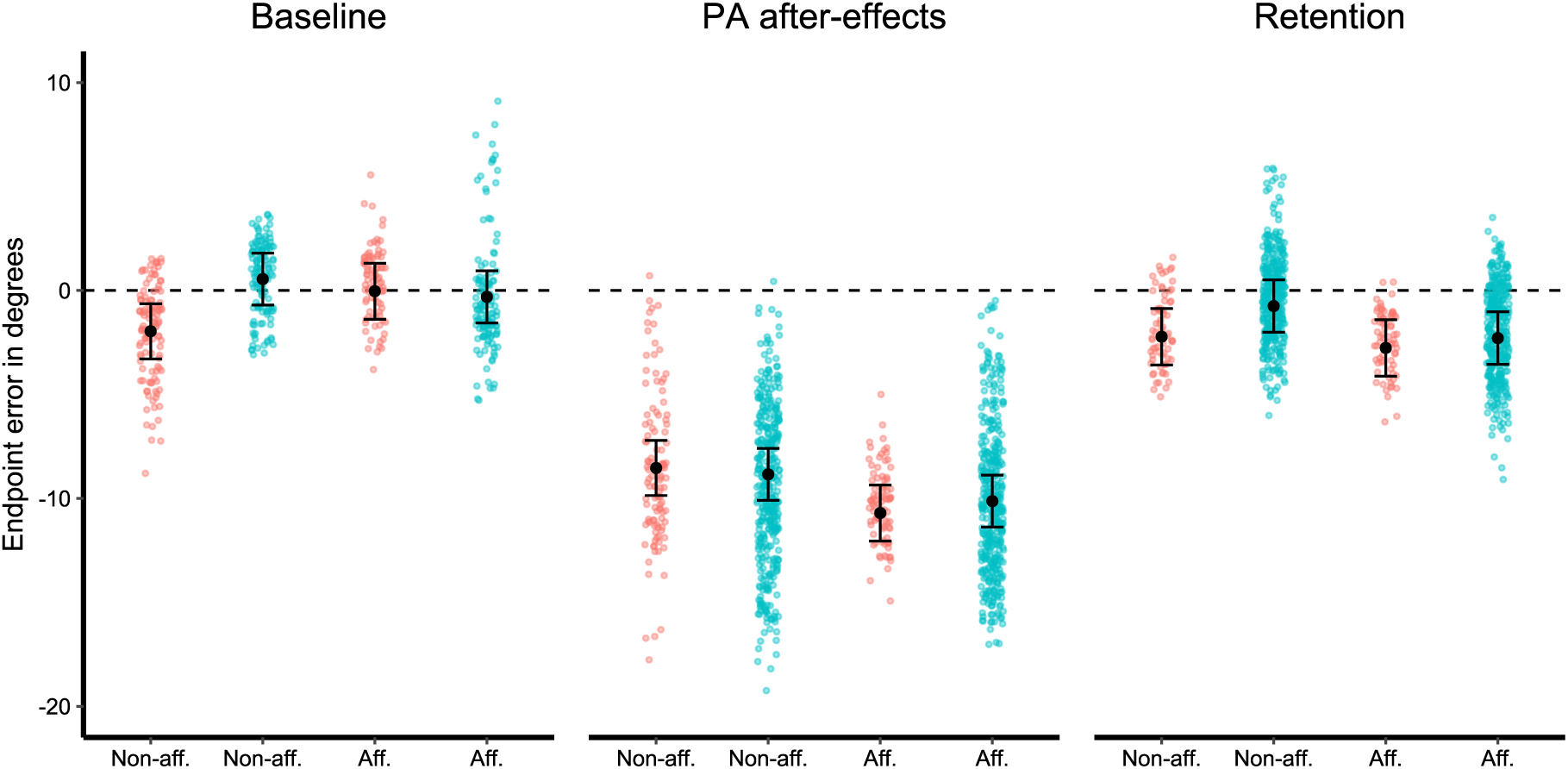
Open-loop endpoint errors for the baseline (B-OL), after-effect (AE-OL), and retention (R-OL) phases are presented for people with CRPS for each hand, split by previous experience with prism adaptation. Eight of the participants with CRPS (red) had previous experience with prism adaptation from a two-week intervention (Halicka, Vittersø, et al., 2020b) ≥ 7 months prior to taking part in the present study. The remaining nine people with CRPS (blue) had no previous experience with prism adaptation, as they either received sham treatment or were not enrolled in the trial. However, not all participants were able to complete prism adaptation with both hands, and data from four participants was removed from the retention phase as they had completed an additional 42 washout trials, resulting in different sample sizes for the baseline and prism exposure phases (previous prism experience: n_affected_ = 6, n_non-affected_ = 8; no experience/sham: n_affected_ = 9, n_non-affected_ = 9) than the retention phase (previous prism experience: n_affected_ = 5, n_non-affected_ = 5; no experience/sham: n_affected_ = 7, n_non-affected_ = 7). Black points and error bars depict the estimated marginal means and the error bars show 95% confidence intervals. The black dashed line shows the target orientation (i.e. zero degree error). Negative values indicate endpoint errors made towards the affected/non-dominant side. Aff. = affected hand; Non-aff. = non-affected hand.

### 6.3 Supplementary Text T1: Summary statistics

#### 6.3.1 Peak velocity

People with CRPS did not differ from controls in the speed at which they performed pointing movements, although there was a tendency for people with CRPS to move slower when using their affected limb. That is, there was no evidence of a difference between Groups (CRPS *M* = 1533.7 mm/s, *SD* = 343.9; controls *M* = 1575.9 mm/s, *SD* = 467.0), or an effect of Hand (affected/non-dominant *M* = 1545.4 mm/s, *SD* = 413.8; non-affected/dominant *M* = 1563.4 mm/s, *SD* = 412.4) on peak velocity averaged across all open-loop and closed-loop trials, *F*s(1, 30) ≤ 0.60, *p*s ≥ .443, ƞ^2^_p_ ≤ .02. There was, however, a significant interaction between Group and Hand, *F*(1, 30) = 9.05, *p* = .005, ƞ^2^_p_ = .23. This interaction appeared to be driven by a smaller peak velocity between the affected side (*M* = 1472.6 mm/s, *SD* = 344.8) and the non-affected side (*M* = 1588.0 mm/s, *SD* = 334.0) for people with CRPS, whereas there was less of a difference between peak velocity for the non-dominant (*M* = 1608.2 mm/s, *SD* = 455.9) and dominant (*M* = 1543.2 mm/s, *SD* = 475.9) hands for controls. None of the differences between conditions was significant after correcting for multiple comparisons, *t*s(30) ≤ 2.36, *p*s_adjusted_ ≥ .100, *d*s ≤ 0.84.

#### 6.3.2 Peak acceleration

Peak acceleration was comparable between people with CRPS and controls, although it was lower when people with CRPS used their affected hand. That is, there were no significant differences between Groups (CRPS *M* = 1058.2 mm/s^2^, *SD* = 3995.8; controls *M* = 10968.1 mm/s^2^, *SD* = 6002.6), or between the Hand used (affected/non-dominant *M* = 10437.8 mm/s^2^, *SD* = 5046.2; non-affected/dominant *M* = 11111.7 mm/s^2^, *SD* = 5212.7) on peak acceleration averaged across all trials, *F*s(1, 30) ≤ 2.55, *p*s ≥ .121, ƞ^2^_p_ ≤ .08. There was a significant interaction between Group and Hand, *F*(1, 30) = 5.99, *p* = .021, ƞ^2^_p_ = .17. Follow-up analyses suggested that this interaction was driven by people with CRPS having lower peak acceleration when using the affected Hand (*M* = 9648.1 mm/s^2^, *SD* = 3683.7) and the non-affected Hand (*M* = 11412.9 mm/s^2^, *SD* = 4078.6), *t*(30) = 2.70, *p*_adjusted_ = .044, *d* = 0.99. There were no significant difference in peak acceleration between the non-dominant (*M* = 11118.0 mm/s^2^, *SD* = 5891.8) and dominant (*M* = 10816.5 mm/s^2^, *SD* = 6109.6) Hand for controls, and or any differences between Groups that depended on the Hand, *t*s(30) ≤ 1.10, *p*s_adjusted_ ≥ .837, *d*s ≤ 0.40.

#### 6.3.3 Baseline closed-loop pointing errors

Endpoint errors during closed-loop pointing were comparable between people with CRPS and controls. There were no significant main effects of Group (CRPS *M* = −0.01°, *SD* =0.43; controls *M* = −0.05°, *SD* = 0.39), or Hand (affected/non-dominant *M* = −0.12°, *SD* = 0.38; non-affected/dominant *M* = 0.06°, *SD* =0.41), or any interactions on endpoint errors for baseline closed-loop trials, *F*s(1, 31) ≤ 2.69, *p*s ≥ .111, ƞ^2^_p_ ≤ .08.

### 6.4 Supplementary Text T2: Model fit

#### 6.4.1 Exponential decay of endpoint errors during prism exposure

Before analysing the constants derived from the fitted models, we analysed the model fit. The model failed to converge, or there was no exponential fit for one person with CRPS (non-affected hand), and for one control (dominant hand). Next, we compared the model fit parameters between Groups and Hand, for those cases where the model did converge and there was an exponential decay (CRPS n = 14; controls n = 17). The results suggested that the models were not found to differ across Groups and Hand. That is, the prediction errors (i.e. the root-mean-square error [RMSE]) which indicated the mean distance from a predicted value to an observed value, for individually fitted models was not significantly different between Groups, (CRPS *M*_RMSE_ = 1.20, *SD* = 0.67; controls *M*_RMSE_ = 0.98, *SD* = 0.72), Hand (affected/non-dominant *M*_RMSE_ = 1.09, *SD* = 0.57; non-affected/dominant *M*_RMSE_ = 1.08, *SD* =0.82), and there was no significant interaction between the two variables, *F*s(1, 29) ≤ 1.70, *p*s ≥ .202, ƞ^2^_p_ ≤ .06. Similarly, there were no significant differences in how much variance was explained by the models (i.e. the *adj. R^2^*) between Groups (CRPS *M_adj.R2_* = .47, *SD* = .26; controls *M_adj.R2_* = .54, *SD* = .26), Hand (affected/non-dominant *M_adj.R2_* = .52, *SD* = .25; non-affected/dominant *M_adj.R2_* = .50, *SD* = .27), and there was no significant interaction between the two variables, *F*s(1, 29) ≤ 0.52, *p*s ≥ .475, ƞ^2^_p_ ≤ .02. As there was no clear difference in the model fits between people with CRPS and controls, or any clear differences depending on the hand used, we proceeded to analyse the constants derived from the models (i.e. 1/*b*, and *c*; Fig. 4).

#### 6.4.2 Exponential decay of endpoint errors during washout

Prior to analysing the constants from the exponential decay function, we analysed the model fit. We were unable to fit an exponential decay function for two participants with CRPS, both for their affected hand. For those cases where the model did converge and there was an exponential decay (CRPS n = 13; controls n = 18), we compared the model fit parameters between Groups and Hand. The prediction error (i.e. RMSE) did not differ between people with CRPS and controls, or for either hand. That is, there was no significant main effect of Group (CRPS *M*_RMSE_ = 0.77, *SD* = 0.35; controls *M*_RMSE_ = 0.59, *SD* = 0.29), Hand (affected/non-dominant *M*_RMSE_ = 0.64, *SD* = 0.27; non-affected/dominant *M*_RMSE_ = 0.69, *SD* = 0.37), and no significant interactions on RMSE, *F*s(1, 29) ≤ 2.03, *p*s ≥ .165, ƞ^2^_p_ ≤ .07. These results suggest that there was no difference in the prediction error between Groups or the hand used. We then analysed how much variance was explained by the models (i.e. the *adj. R^2^*). There were no significant differences between Groups (CRPS *M_adj.R2_* = .34, *SD* = .23; controls *M_adj.R2_* = .42, *SD* = .30), *F*(1, 29) = 0.43, *p* = .517, ƞ^2^_p_ = .01. There was a tendency for models to explain a greater proportion of the variance for models fitted to data from the non-affected/dominant hand (*M_adj.R2_* = .41, *SD* = .27) than the affected/non-dominant hand (*M_adj.R2_* = .35, *SD* = .28), although not statically significant, *F*(1, 29) = 3.16, *p* = .086, ƞ^2^_p_ = .10. Neither did we find any evidence that this tendency varied between Groups, as there was no significant interaction between Group and Hand on *adj. R^2^*, *F*s(1, 29) ≤ 1.14, *p*s ≥ .343, ƞ^2^_p_ ≤ .04. Therefore, as there was no clear difference in the model fits between people with CRPS and controls, and the tendency for the models to explain a greater proportion of the variance for the non-affected/dominant hand did not vary between groups, we proceeded to analyse the constants derived from the models (i.e. 1/*b*, and *c*; Fig. 5).

#### 6.4.3 Feedforward motor control

Prior to analysing the detrended data for initial trajectory orientations, we inspected the model fit. We were unable to fit initial trajectory orientations to the exponential decay function for one person with CRPS (non-affected hand), and two controls (one non-dominant hand, one dominant hand). For those cases where we were able to fit an exponential decay to their initial trajectory orientation, we compared the model fit parameters between Groups and Hand. In general, the models fitted to the initial trajectory orientations had greater prediction error (*M*_RMSE_ = 10.58, *SD* = 4.72) and explained less of the variance (*M_adj.R2_* = .02, *SD* = .06) than the models fitted to endpoint errors (*M*_RMSE_ = 1.09, *SD* = 0.70; *M_adj.R2_* = .51, *SD* = .26), which indicates that the exponential decay was a better fit for endpoint errors than for initial trajectory orientations. For initial trajectory orientations there was a tendency for the prediction error of the model to be greater for controls (*M*_RMSE_ = 11.80, *SD* = 5.90) than people with CRPS (*M*_RMSE_ = 9.24, *SD* = 2.40), although not significant, *F*(1, 28) = 3.19, *p* = .085, ƞ^2^_p_ = .10. There was no significant difference in RMSE between the Hand used (affected/non-dominant *M*_RMSE_ = 10.10, *SD* = 3.42; non-affected/dominant *M*_RMSE_ = 11.08, *SD* = 5.83), and there was no significant interaction with Group and Hand, *F*s(1, 28) ≤ 0.80, *p*s ≥ .379, ƞ^2^_p_ ≤ .03. This suggests that the prediction error did not vary depending on the hand used, although there was a trend for models to make greater prediction errors for controls participants than people with CRPS. Next we analysed how much of the variance in the data was accounted for by the models (i.e. *adj. R^2^*). There were no significant differences between Groups (CRPS *M_adj.R2_* = .04, *SD* = .08; controls *M_adj.R2_* = .01, *SD* = .04), or Hand (affected/non-dominant *M_adj.R2_* = .03, *SD* = .07; non-affected/dominant *M_adj.R2_* = .02, *SD* = .06) on *adj. R^2^*, and no significant interaction, *F*s(1, 28) ≤ 2.39, *p*s ≥ .134, ƞ^2^_p_ ≤ .08. This suggests that the amount of variance explained by the models did not differ between people with CRPS and controls, or depending on the Hand used. However, it should be noted that the amount of variance explained by an exponential fit for initial trajectory orientations were substantially lower than that of endpoint errors.

### 6.5 Supplementary Text T3: Exploratory analysis – Covariate analyses

People with unilateral CRPS have been reported to have bilateral proprioceptive deficits (Bank et al., 2013). In previous research, absolute pointing errors made with the unseen hand(s) have been interpreted as evidence of deficits in arm position sense in people with CRPS (Lewis et al., 2010). Therefore, we used the absolute endpoint error during the baseline Open-loop Block (“Absolute Baseline Error”) as a proxy measure of proprioceptive accuracy. Our results, however suggest that there were no differences in Absolute Baseline Error between people with CRPS and controls, *F*s(1, 28) ≤ 0.06, *p*s ≥ .805, ƞ^2^_p_ < .01. When we re-ran our primary analysis (3.3.1.) of open-loop endpoint errors using ANCOVAs with Absolute Baseline Error included as a covariate, we found that it did not influence our results, *F*s(2, 62) ≤ 1.03, *p*s ≥ .362, ƞ^2^_p_ ≤ .03. Next, because more of our participants with CRPS than controls were exposed to leftward-shifting prisms, we considered that the direction of the prismatic shift might have influenced the degree of sensorimotor realignment, as a difference between adapting to leftward and rightward shifting lenses has been reported previously (Redding & Wallace, 2009). We also did not observe any influence of the direction of the prismatic shift on our results from follow-up ANCOVAs, *F*s(2, 62) ≤ 0.62, *p*s ≥ .544, ƞ^2^_p_ ≤ .02, which suggests that our findings are unlikely to be due to a greater number of people with CRPS being exposed to leftward shifting prisms than controls. Therefore, it does not seem likely that our findings can be attributed to differences in proprioceptive abilities, the direction of the prismatic shift, and/or the counterbalancing order.

We considered that the counterbalancing order might influence our results, as inter-limb transfer has previously been found to be greater when the dominant hand is adapted first, compared to the non-dominant hand (Redding & Wallace, 2008, 2011). When we reanalysed the endpoint errors with Counterbalancing Order (affected/non-dominant first, non-affected/dominant first), Hand, and Open-loop Block as independent variables. There was no significant main effect of Counterbalancing Order on endpoint errors, and/or no significant interactions with Hand, or Open-loop Block, *Fs*(2, 62) ≤ 2.64, *p*s ≥ .080, ƞ^2^_p_ ≤ .08. These results therefore suggest that there was no significant influence of inter-limb transfer on endpoint errors.

### 6.6 Supplementary Text T4: Exploratory analysis – Kinematic changes

As we anticipated the kinematic data to be noisy, we first analysed the strength of individual correlations for each Hand, pooling the data for each Group. That is, we compared *t*-values to zero for each Hand, averaged across all participants. The t values for participants’ non-affected/dominant hand significantly deviated from 0, indicating the presence of a linear relationship between endpoint errors (trial_n_) and the subsequent change in initial trajectory orientation (trial_n+1_) (*M_t_* = −0.46, *SD* = 1.23), *t*(32) = 2.15, *p* = .039, *d* = 0.37. This association suggests that when participants made endpoint errors in a given direction they adjusted their movement plan in the opposite direction on the subsequent trial (e.g. if an endpoint error was made towards the right, the subsequent movement was angled more towards the left). We did not observe evidence of such a linear relationship when people used their affected/non-dominant hand (*M_t_* = −0.18, *SD* = 1.24), *t*(31) = 0.83, *p* = .416, *d* = 0.15. Therefore, the data pooled across all participants provides evidence that feedforward motor control was used to reduce trial-by-trial endpoint errors relative to the previous trial for the non-affected/dominant hand - but not the affected/non-dominant hand - during early prism exposure trials.

Unilateral Kolmogorov-Smirnov distribution tests (Vindras et al., 2012) on the *p*-values obtained from individual correlations analyzed pooled across Groups, reveled no evidence that *p*-values were biased towards zero for either hand, *D^+^* ≤ 0.18, *ps* ≥ .104. These findings suggest that the *p*-values for the correlation between endpoint errors for early trials (*n*) and the change in initial trajectory orientation on the next trial (i.e. trial*_n_*_+1_ -trial*_n_*), were not biased towards zero.

Taken together, there was evidence that feedforward motor control was used to reduce endpoint errors during early prism exposure when participants used the non-affected/dominant hands, but only from the analysis of *t*-values. There was no evidence of such use of feedforward motor control from *t*-values when participants used the affected/non-dominant hands. Therefore, the evidence of feedforward motor control during early trials for the non-affected/dominant hands should be interpreted with caution.

## Footnotes

1 CRPS, Complex Regional Pain Syndrome.

